## Extended Data for "Mapping mechanical stress in curved epithelia of designed size and shape"

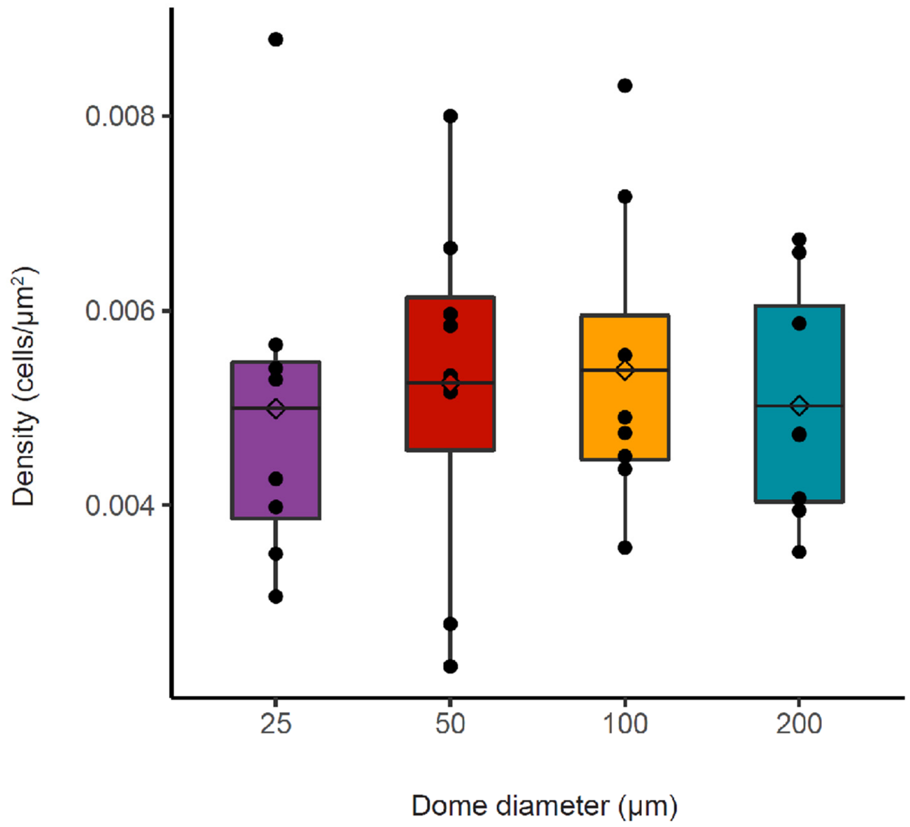


**Extended Data Figure 1 | Dome cell density as function of footprint diameter.**
Density of cells in MDCK spherical domes with 25, 50, 100 and 200 µm footprint diameter. Data are shown as median ± SD of n=8 (all conditions conditions).

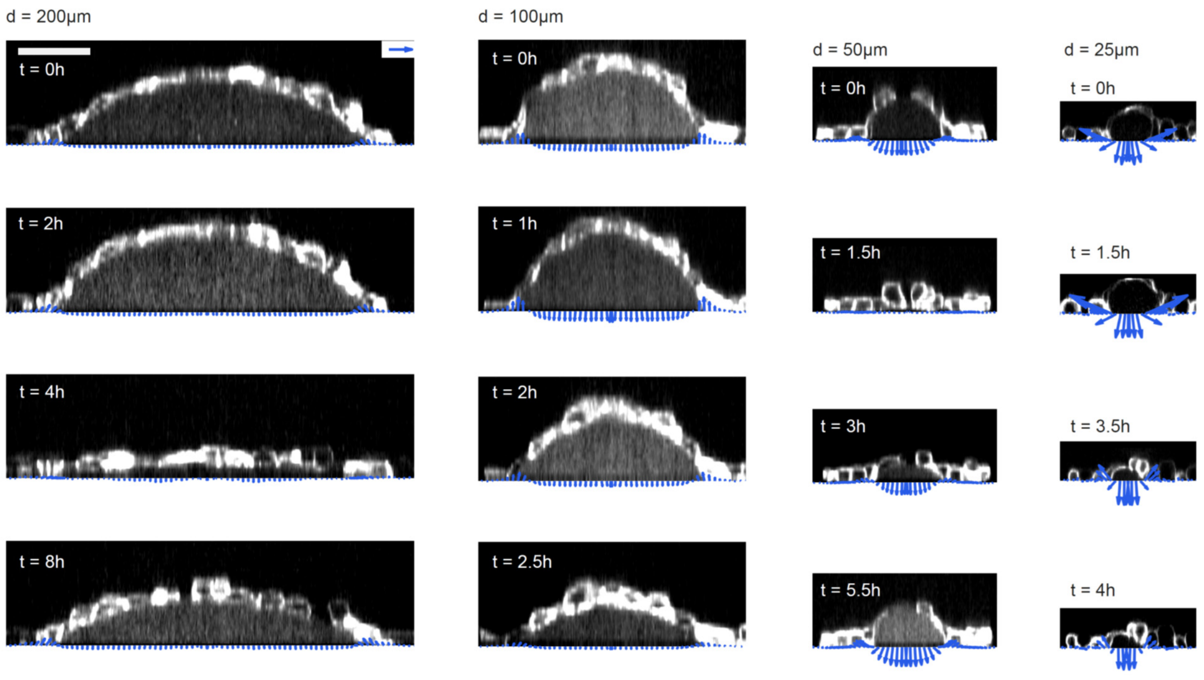


**Extended Data Figure 2 | Time evolution of spherical domes of different sizes.**
Time evolution of radially-averaged tractions on lateral views of domes of 200, 100, 50 and 25 µm diameter footprints (from left to right). Blue vectors represent the radial and vertical components of the tractions. Scale bar, 50µm. Scale vector, 100 Pa.

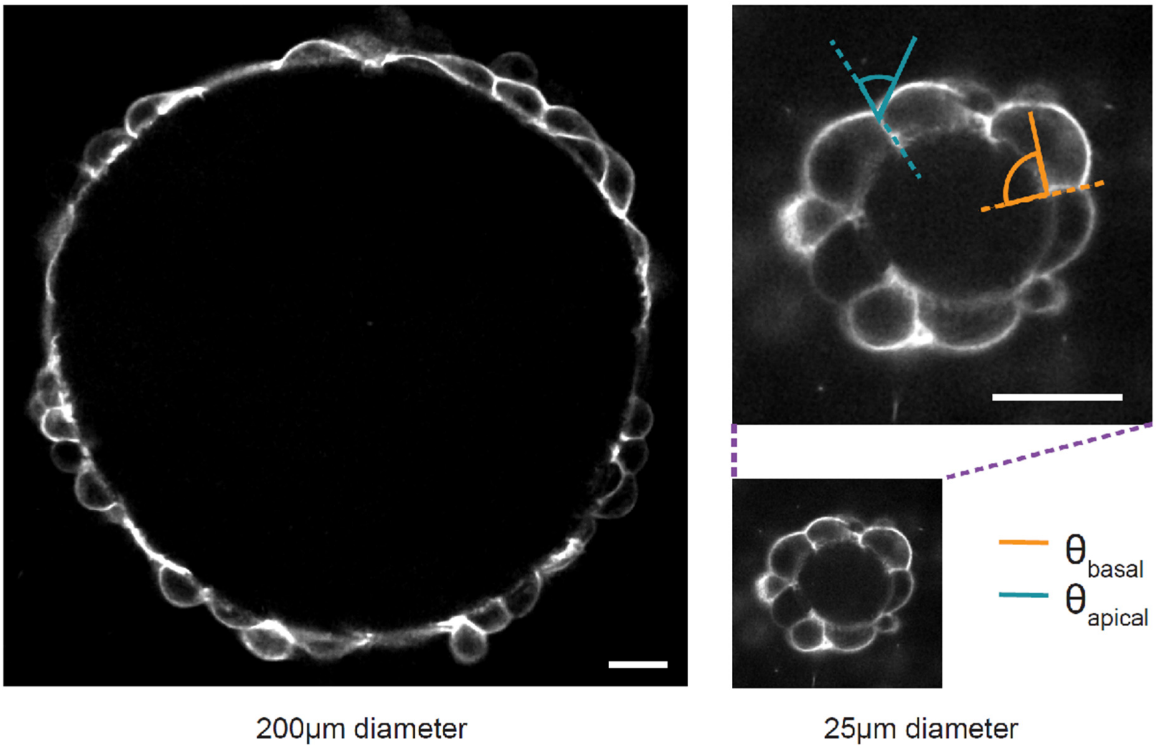


**Extended Data Figure 3 | Contact angle between cells in domes.**
Confocal slices of 200 μm (left) and 25μm (right) patterned-diameter domes showing the apical (blue) and basal (orange) contact angles between cells. Scale bar: 20μm.

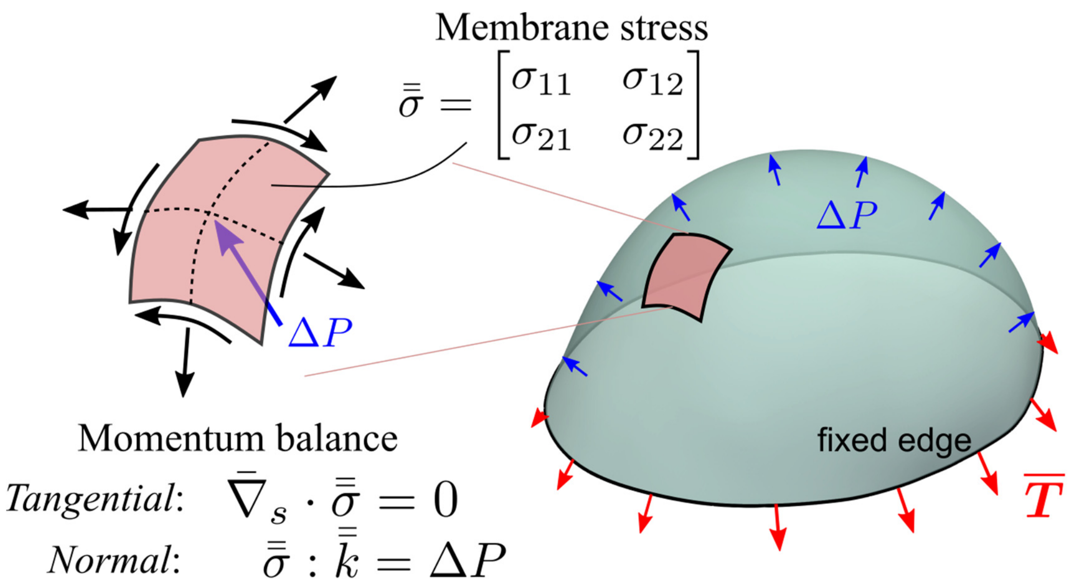


**Extended Data Figure 4 | Schematic illustrating cMSM**

The monolayer stress tensor $\bar{\bar{\sigma}}$ is inferred from luminal pressure $\Delta P$ and monolayer shape, solving the two tangential equilibrium equations and the out-of-plane force balance. As a consequence of luminal pressure, the substrate exerts a traction $\vec{T}$ on the free-standing monolayer (note that $\vec{T}$ shown here has the opposite sign than that reported in the main text, which indicates the traction generated by the cells on the substrate).

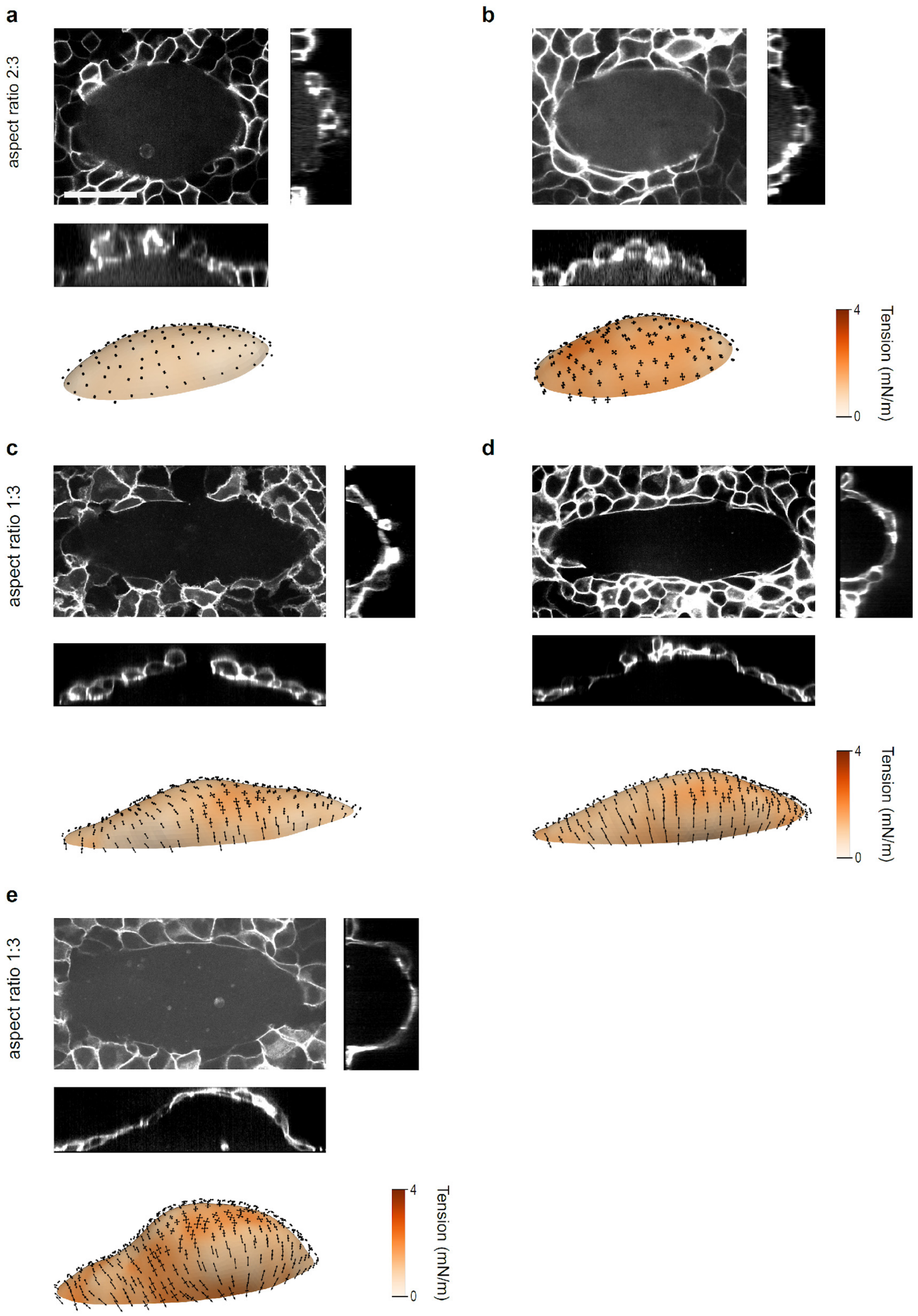


**Extended Data Figure 5 | Examples of stress maps on ellipsoidal caps**
Each panel shows a top view and two lateral views of the ellipse and the stress reconstruction below.

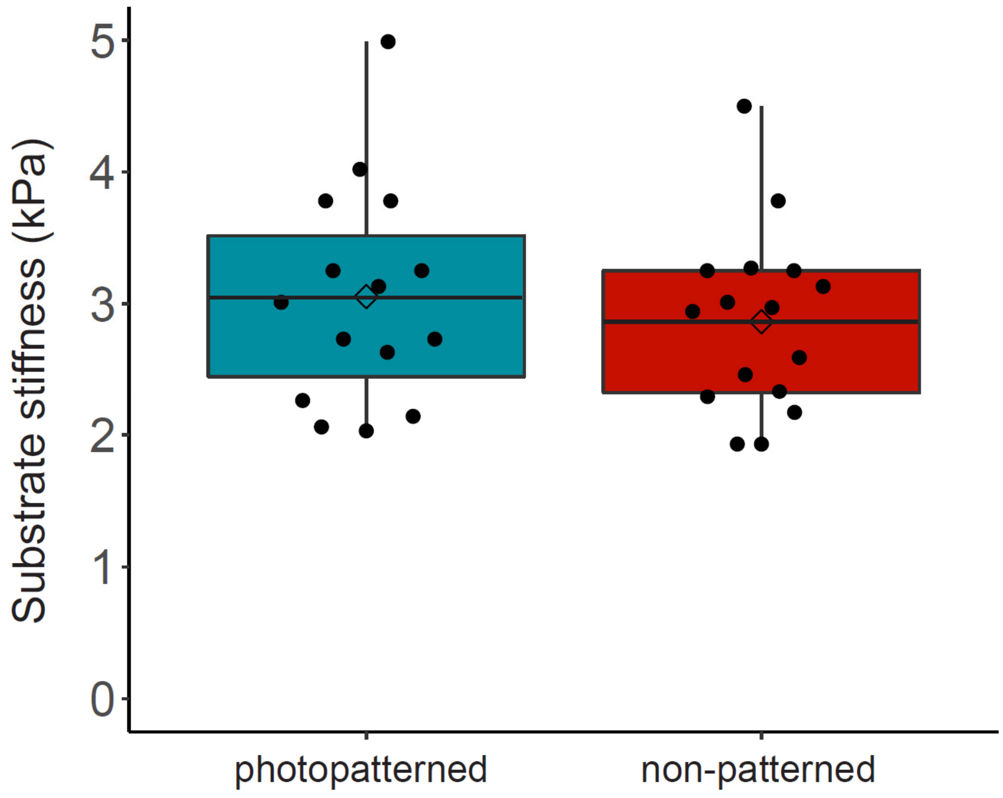


**Extended Data Figure 6 | Stiffness of the gel substrate before and after photopatterning.** Gel stiffness was measured using the ball indentation method in photopatterned (n=15) and non-patterned (n=16) gels. The process of patterning did not affect substrate stiffness significantly (*P*-value = 0.58).
