## Supplementary notes for "Mapping mechanical stress in curved epithelia of designed size and shape"

### Supplementary Materials for *Mapping mechanical stress in curved epithelia of designed size and shape*

### Supplementary Note 1

### Computational vertex model with curved surfaces for 3D epithelia

To study the mechanics of epithelial domes, we implement a 3D computational vertex model with curved cell surfaces. We adopt a conventional approach according to which cell surfaces experience constant and possibly non-uniform surface tension representing contractility of the actomyosin cortex and cell volumes are fixed [1].

#### 1.1 Polygonal tessellation

The epithelial cells within a monolayer are assumed to be polyhedra composed of polygonal apical and basal faces, and rectangular lateral faces. To allow for curved surfaces, each face is further discretized using linear triangular elements (Supplementary Fig. 1a,b). Cells are then assembled by tying together the shared edges across contiguous cell faces through equality constraints [2]. Following our observations, we further assume that during dome inflation, no topological rearrangements occur within the tissue. Cells retain their original connectivity and tissue deformation is only accommodated through cell shape changes. Assembling a monolayer from the cells is therefore achieved by stitching together the contiguous lateral faces between cells through additional equality constraints. Through these physical constraints, the lateral cell surfaces of neighboring cells form cell-cell contacts.

#### 1.2 Surface tension generated by the actomyosin cortex

We assume that the actomyosin cortex, which lines the interior of each cell face, is the main subcellular element determining epithelial mechanics. Due to the longer timescale of dome inflation compared to actomyosin dynamics, we assume that all cortical dynamics are at steady-state during dome inflation and that the cortex generates a constant and isotropic active surface tension $\gamma$, which can be different on each cell face (Supplementary Fig. 1a). We therefore express the mechanics of the actomyosin cortex on a face $f$ in the tissue using a conventional virtual work function

$$\delta W_f(\boldsymbol{x}_1, \ldots, \boldsymbol{x}_{N_f}) = \gamma_f \delta A_f(\boldsymbol{x}_1, \ldots, \boldsymbol{x}_{N_f}) \tag{1}$$

where $\boldsymbol{x}_i$ denotes the position of node $i$ in the triangulation of face $f$, $\gamma_f$ is the active tension generated by the actomyosin cortex and $A_f$ the corresponding face surface area. We assumed that apical and basal tension is contractile, and that contractility on the lateral faces dominates over adhesion, so that $\gamma_f > 0$ for all faces. For simplicity, we did not consider the effects of actin cables or other complex cortical architectures on surface tension.

#### 1.3 Volume conservation

Following previous observations [3] and to avoid cell collapse due to contraction of all faces, we assume each individual cell volume $\Omega_c$ to be conserved during cellular deformations. Volume conservation is imposed through a Lagrange multiplier requiring that the change in cell volume remains zero

$$\Omega_c(\boldsymbol{x}_1, \ldots, \boldsymbol{x}_{N_c}) - \Omega_c^0 = 0, \tag{2}$$

where $\Omega_c^0$ is the initial volume of cell $c$ and $N_c$ the number of nodes on cell $c$.

#### 1.4 Dome inflation

We further define an adherent region of the basal surface to the substrate in which nodes' movements are restricted and allow for nodes in a non-adherent region to move freely. The volume of the lumen enclosed between the non-adherent region and the substrate is then incrementally increased through a Lagrange multiplier requiring the lumen volume at each step of the dome inflation to be equal to the imposed volume

$$\Omega_l(\boldsymbol{x}_1, \ldots, \boldsymbol{x}_{N_l}) = \Omega^*, \tag{3}$$

where $\Omega_c$ is the lumen volume calculated using nodes positions and $N_l$ the number of nodes that define the surface bounding the lumen.

#### 1.5 Lagrangian of a tissue

The Lagrangian of the tissue is simply built by taking into account the virtual work function and the constraints of the problem for each cell and face of the tissue

$$\delta\mathcal{L} = \sum_f \gamma_f \delta A_f - \sum_c \Delta P_c \left(\delta\Omega_c - \delta\Omega_c^0\right) + \Delta P_l(\delta\Omega_l - \delta\Omega^*), \tag{4}$$

where the summations are performed over all faces and cells in the tissue. The Lagrange multiplier $\Delta P_c$ is the pressure difference across the cell interior and the exterior medium. The Lagrange multiplier $\Delta P_l$ gives a direct readout of the pressure inside the dome lumen (Supplementary Fig. 1a,c). For arbitrary virtual vertex displacements, virtual cell pressure changes and virtual lumen pressure changes; and by making the Lagrangian stationary, Eqs. (4) provide a system of coupled nonlinear equations enforcing mechanical equilibrium of the network nodes, along with cell volume and lumen volume constraints. However, as discussed in [4], solving Eqs. (4) leads to uncontrolled mesh distortion as nodes can move tangentially without changing cell area or volume. To avoid such tangential motions of nodes, we added an effective cortical viscosity, which vanishes at equilibrium to avoid biasing the end results. Therefore, the dome inflation process can be seen as the quasi-static evolution of active-viscous cortices with fixed cellular volume and an increasing lumen volume.

#### 1.6 Measuring tension using Young-Laplace's law

For spherical epithelial domes, we use Young-Laplace's law to estimate tension within the monolayer. At each time-step, we compute the dome radius $R$ from the tissue height at the apical-basal mid-plane $h$ and the basal footprint radius $R_b$ (Supplementary Fig. 1b,d)

$$R = \frac{h^2 + R_b^2}{2h}. \tag{5}$$

Tension is then obtained from $R$ and $\Delta P_l$ as

$$\sigma = \frac{R\Delta P_l}{2}. \tag{6}$$

The areal strain $\varepsilon$ of the dome shown was computed as

$$\epsilon = \frac{\pi(R_b^2 + h^2)}{\pi R_b^2} - 1 = \frac{h^2}{R_b^2}, \tag{7}$$

where $\pi(R_b^2 + h^2)$ is the surface area dome and $\pi R_b^2$ the surface area of the basal footprint.

#### 1.7 Tissue constitutive relation

As shown in [5], the tension-strain relationship of domes can be captured by the tension-strain relation of an idealised tissue model. Considering the idealised tissue made of identical cells as regular hexagonal prisms with constant volume subjected to uniform equibiaxial stretch, the tissue surface tension is expressed through the following cellular constitutive relation [5]

$$\sigma = \gamma_a + \gamma_b - \gamma_l \frac{k}{(\varepsilon_c + 1)^{3/2}}, \tag{8}$$

where $\sigma$ is the tissue effective surface tension, $\gamma_a$ the apical surface tension, $\gamma_b$ the basal surface tension, $\gamma_l$ the lateral surface tension, $\varepsilon_c$ is the tissue areal strain and $k$ a non-dimensional geometrical constant. As the tissue stretches, contribution to tissue tension from the lateral faces decreases, therefore tissue tension should saturate to the apico-basal surface tensions.

#### 1.8 Tensional asymmetry

Given our observation that the contact angle between adjacent cells was smaller on the apical surface compared to the basal surface, we concluded that surface tension on the apical side was smaller compared to basal surface tension, which could give rise to bending moments affecting the mechanical balance of domes. More specifically, in addition to tension, bending moments could contribute to balance the pressure difference across the tissue. Such tensional asymmetry has been shown to produce spontaneous curls on free edges of flat free-standing cell monolayers [6], thereby endowing the tissue with an effective spontaneous curvature. Since spontaneous curvature induces a length-scale, we reasoned that these bending moments should manifest differently for domes of different sizes. To investigate such hypothesis, we first inflated domes of different circular footprint diameters up to a 100% areal strain (Supplementary Fig. 2a-f) at a high apical-to-basal surface tension ratio (1:9 ratio) and calculated tissue tension as a function of dome areal strain using Young-Laplace's law 6. Our simulations showed that the tension-strain curves varied little between the different domes sizes throughout the inflation process (Supplementary Fig. 2g) and were close to the analytical estimate for a planar and uniform tissue given by Eq. (8), which is independent of tension asymmetry. Furthermore, all these curves were compatible with our experimental observations (Fig. 1.e in the main text).

To further assess the relevance of surface tension asymmetry on the tension-strain curve, we calculated the tissue surface tension using Laplace's law for large domes inflated at various apical-basal surface tension degrees of asymmetry, with tension ratios ranging between 1:9 to 1:1 while keeping the average apico-basal tension fixed (Supplementary Fig. 2h), therefore varying the strength of bending moments. We found that all tension-strain curves were very close during the entire inflation process, and also closely followed the curve given by Eq. (8). We thus concluded that tension asymmetry, tissue spontaneous curvature or bending moments did not have a substantial mechanical effect in balancing the transmural pressure, and hence do not need to be included in the interpretation of epithelial bulge tests to compute tissue tension.

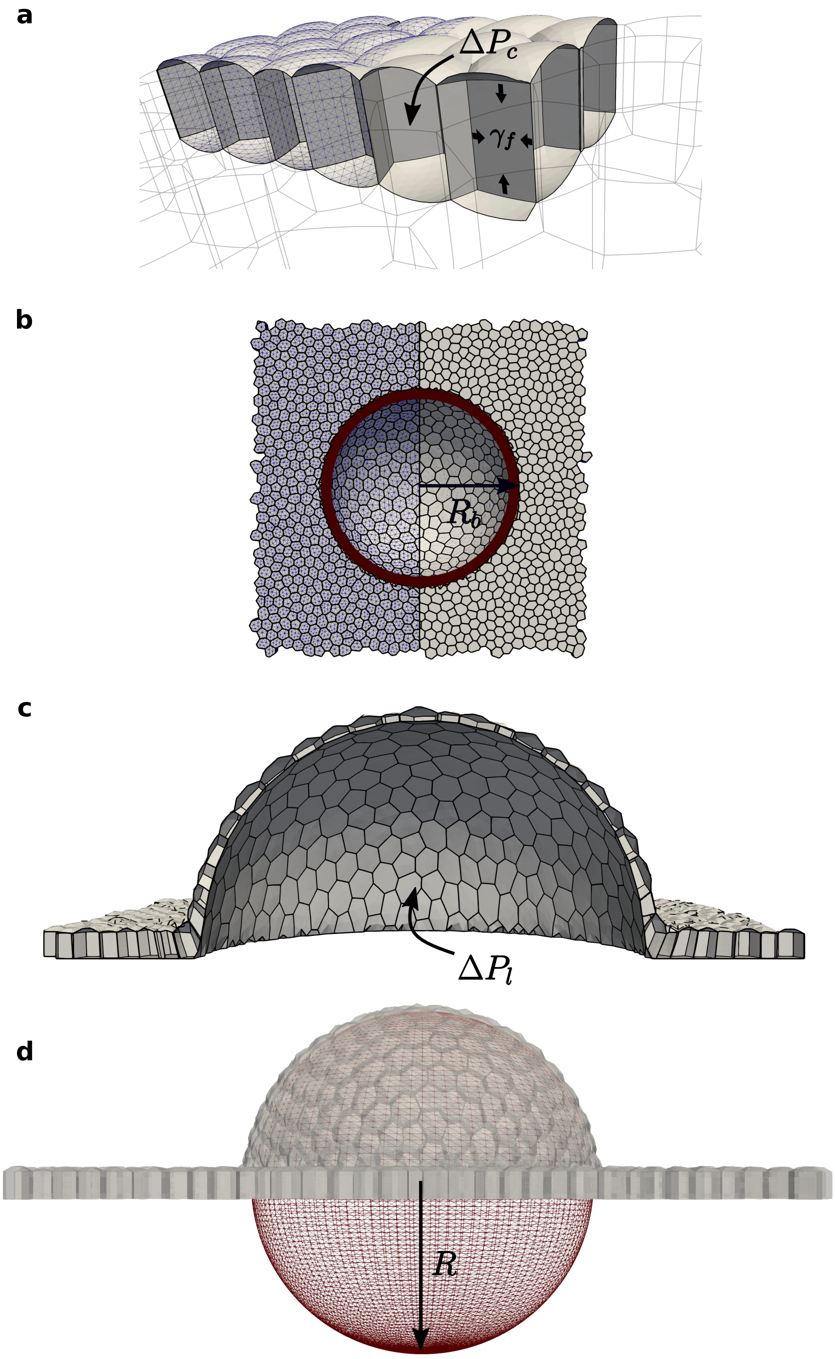

Supplementary Figure 1: Computational vertex model with curved surfaces. **(a)** Constant and isotropic surface tension on each face $\gamma_f$ along with cell volume conservation generating a cellular pressure that bulges non-adhered surfaces outwards. **(b)** We defined a non-adherent region of radius $R_b$ on the basal footprint outside which nodes are fixed. **(c)** We imposed an incremental volume increase in the non-adherent region $\Omega_l$ through a Lagrangian multiplier providing a direct readout of lumen pressure $\Delta P_l$ involved in Laplace's law. **(d)** Inflated domes can be closely approximated as spherical caps.

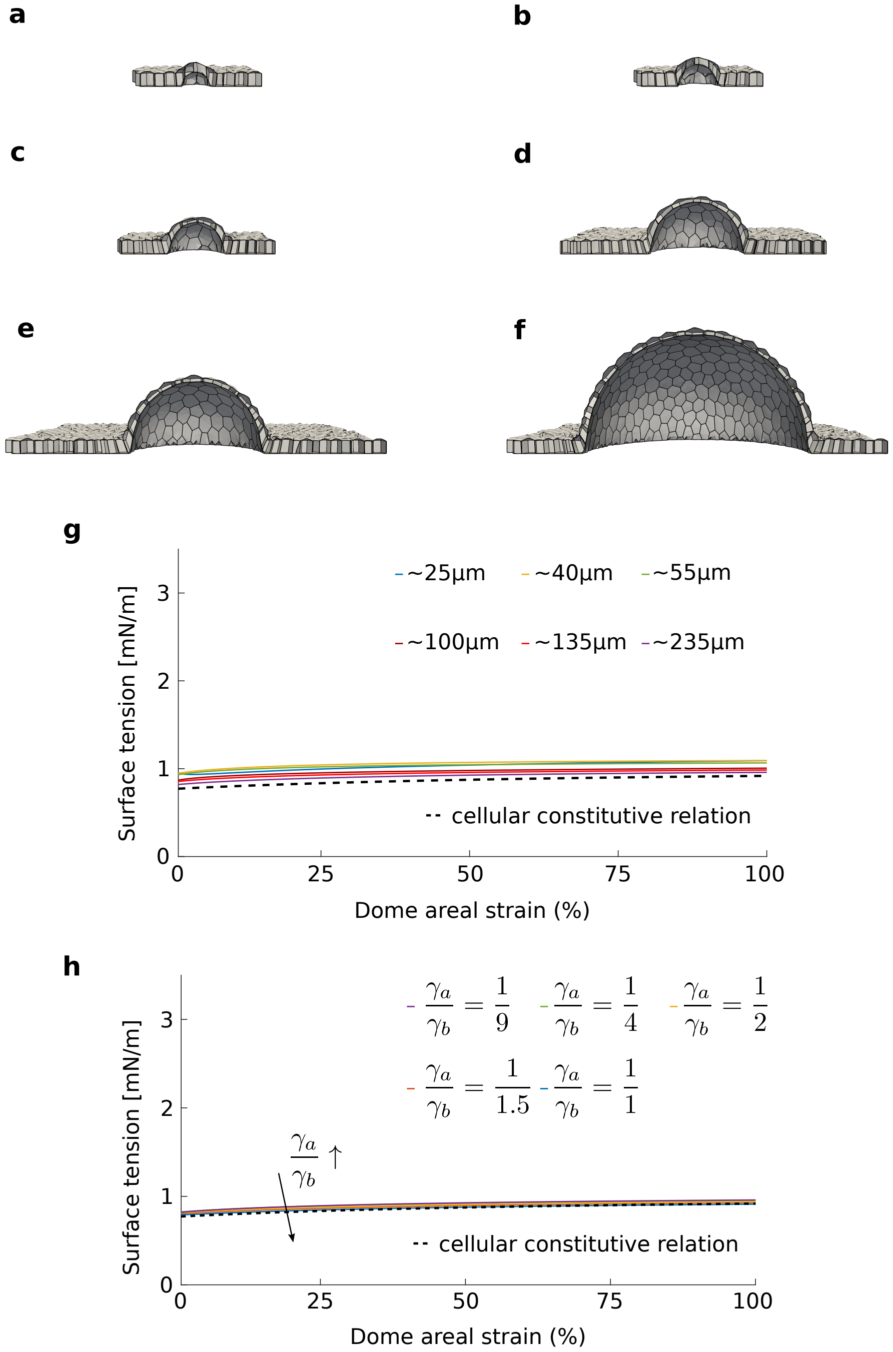

Supplementary Figure 2: Surface tension measured in computational vertex model simulations using Laplace's law show negligible differences between different dome sizes and surface tension asymmetries. **(a-f)** Cross-sections of domes with different basal footprint sizes at 100% areal strain and 1:9 apical-to-basal surface tension ratio. **(g)** Surface tension as a function of strain for the different dome basal footprint diameters: $\sim 25\,\mu\mathrm{m}$ (blue), $\sim 40\,\mu\mathrm{m}$ (yellow), $\sim 57\,\mu\mathrm{m}$ (green), $\sim 100\,\mu\mathrm{m}$ (brown), $\sim 140\,\mu\mathrm{m}$ (red), $\sim 235\,\mu\mathrm{m}$ (purple). The black dashed curve shows the cellular constitutive relation shown in Eq. (8) for equivalent cellular areal strains. **(h)** Surface tension as a function of strain at different apical-to-basal surface tension ratios for the dome shown in (**f**). The black dashed curve shows the cellular constitutive equation shown in Eq. (8) for equivalent cellular areal strains. *Parameters:* $\gamma_l = 0.1\,\mathrm{mN\,m^{-1}}$ for all curves*;* $\gamma_a = 0.1\,\mathrm{mN\,m^{-1}}$, $\gamma_b = 0.9\,\mathrm{mN\,m^{-1}}$ (purple)*;* $\gamma_a = 0.2\,\mathrm{mN\,m^{-1}}$, $\gamma_b = 0.8\,\mathrm{mN\,m^{-1}}$ (green)*;* $\gamma_a = 0.333\,\mathrm{mN\,m^{-1}}$, $\gamma_b = 0.666\,\mathrm{mN\,m^{-1}}$ (yellow)*;* $\gamma_a = 0.4\,\mathrm{mN\,m^{-1}}$, $\gamma_b = 0.6\,\mathrm{mN\,m^{-1}}$ (red)*;* $\gamma_a = 0.5\,\mathrm{mN\,m^{-1}}$, $\gamma_b = 0.5\,\mathrm{mN\,m^{-1}}$ (blue).

### Supplementary Note 2

### Curved Monolayer Stress Microscopy (cMSM): inferring surface stresses on inflated membranes

#### 2.1 Balance equations for inflated membranes

We consider a thin interface represented by its midsurface $\Gamma$ embedded in $\mathbb{R}^3$ (Supplementary Fig. 3). The surface is parametrized by the mapping $\boldsymbol{x} = \varphi(\boldsymbol{\xi})$ where $\boldsymbol{\xi} \in \Gamma_0 \subset \mathbb{R}^2$ belongs in the parametric space with Cartesian coordinates $\{\xi^1, \xi^2\}$. The covariant basis vectors tangent to the surface are then defined as [7]

$$\boldsymbol{e}_a = \frac{\partial \boldsymbol{x}}{\partial \xi^a} \tag{9}$$

where $a \in \{1, 2\}$. The first fundamental form of the surface is expressed as

$$g_{ab} = \boldsymbol{e}_a \cdot \boldsymbol{e}_b. \tag{10}$$

The surface normal is a unit vector given as

$$\boldsymbol{n} = \frac{\boldsymbol{e}_1 \times \boldsymbol{e}_2}{|\boldsymbol{e}_1 \times \boldsymbol{e}_2|}. \tag{11}$$

When the thickness of the interface is smaller than other relevant dimensions, membrane theory is applicable. In this theory the wall stresses are assumed to be tangential to $\Gamma$. The membrane state of stress on $\Gamma$ is captured by the surface tension tensor denoted by $\boldsymbol{\sigma} = \sigma^{ab} \boldsymbol{e}_a \otimes \boldsymbol{e}_b$. On an

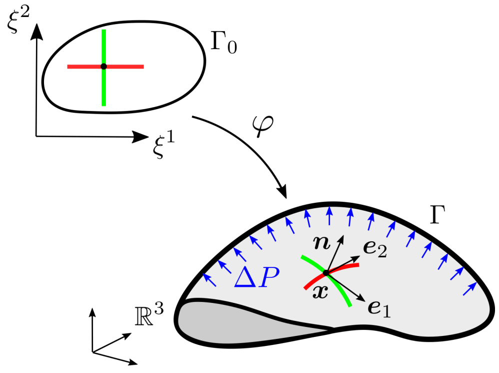

Supplementary Figure 3: Membrane midsurface $\Gamma$ parametrized by the mapping $\boldsymbol{x} = \varphi(\boldsymbol{\xi})$.

open surface $\Gamma$ with a boundary edge $\partial\Gamma$ and edge normal $\boldsymbol{n}_l$ (tangential to the surface), the edge traction vector is given as $\boldsymbol{t} = \boldsymbol{\sigma}\boldsymbol{n}_l$. The balance of angular momentum implies that $\boldsymbol{\sigma}$ is symmetric. The balance of linear momentum in tangential directions in the absence of any tangential body forces is given by [8, 9]

$$\boldsymbol{\nabla}_s \cdot \boldsymbol{\sigma} = 0, \tag{12}$$

where $\boldsymbol{\nabla}_s \cdot$ is the surface divergence operator. The balance of linear momentum in the normal direction is given by the generalized Young-Laplace relation

$$\boldsymbol{\sigma} : \boldsymbol{\kappa} = \Delta P, \tag{13}$$

where $\boldsymbol{\kappa}$ is the second fundamental form of $\Gamma$ and $\Delta P$ is the pressure difference across the interface. Note that the surface divergence operator and $\boldsymbol{\kappa}$ depend on the shape of the surface.

We note that the description of the monolayer as a membrane with a midsurface supporting a state of membrane stress, leading to the above equations, is a fundamental modeling assumption of the stress inference method proposed here. This assumption may be inadequate depending on the system and particularly for smaller domes and for domes at low inflation levels, where midsurface deformations are of the order of monolayer thickness and bending modulus may become relevant. We tested this assumption by analyzing 3D vertex models of finite thickness and exhibiting strong apico-basal tension asymmetry as discussed in Supplementary note 1.

As in 2D, there are infinitely many divergence-free symmetric surface tensor fields on a given deformed shape satisfying Eq. (12). Eq. (13) provides an additional constraint on such tensor fields. Eqs. (12) and (13) constitute a system of three differential-algebraic equations that may provide a statically determinate system to find the three independent components of the symmetric surface tension tensor. To the best of our knowledge, the precise conditions for this problem to be mathematically well-posed that lead to a unique surface tension tensor are not known, but experience with computational approaches (see next section) suggest that these equations enable the determination of the surface tension tensor. For special cases, like simply connected axisymmetric surfaces, the problem can be analytically solved and is known to be well-posed [10, 11].

For an open patch of an inflated membrane, if the edge tension on the boundary $\boldsymbol{t}$ is prescribed, then Eqs. (12) and (13) are augmented with an additional boundary condition $\boldsymbol{t} = \boldsymbol{\sigma}\boldsymbol{n}_l$ that is necessary for static equilibrium. It is unclear whether the boundary data $\boldsymbol{t}$ is strictly required in general for the well-posedness of the stress inference problem on an open surface. In any case, it is clear that these tractions should satisfy a compatibility constraint $\int_{\partial\Gamma} \boldsymbol{t} d\ell = \Delta P \int_{\Gamma} \boldsymbol{n} dS$ coming from global force balance. For special cases such as simply connected axisymmetric surfaces, the edge tensions do not need to be specified as the symmetry of the problem decides the only possible value of edge traction vector that is compatible with the solution.

Next, we introduce previous approaches proposed to solve this inverse problem.

##### 2.1.1 Approaches for stress inference on inflated membranes

We first discuss the solution to the surface tension inference problem on axisymmetric surfaces. Beyond axisymmetry and for generic membrane shapes, Eqs. (12) and (13) need to be solved numerically to determine $\boldsymbol{\sigma}$. We then summarize several computational approaches that have been proposed to address this problem.

##### Closed-form solution for axisymmetric membranes

For axisymmetric membranes, the governing equations are simplified to the following form:

$$\frac{d}{dr}(\sigma_1^1 r) = \sigma_2^2, \tag{14}$$

$$\frac{d}{dr}\sigma_1^2 = -\frac{3\sigma_1^2}{r}, \tag{15}$$

$$\sigma_1^1\kappa_1^1 + \sigma_2^2\kappa_2^2 = \Delta P, \tag{16}$$

where $\kappa_1^1$ and $\kappa_2^2$ are principal curvatures of the surface of revolution in meridian and azimuthal directions and $r$ is the radial distance from the axis of revolution. $\sigma_b^a$ are components of $\boldsymbol{\sigma}$ expressed in the principal directions. Additionally, for axisymmetric surfaces, the principal curvatures are related by

$$\frac{d}{dr}(r\kappa_2^2) = \kappa_1^1. \tag{17}$$

Using this relation, Eq. (14) can be solved directly [10, 11] to obtain

$$\sigma_1^1 = \frac{1}{\kappa_2^2}\left(\frac{\Delta P}{2} + \frac{c_0}{r^2}\right), \tag{18}$$

where $c_0$ is a constant of integration to be determined. If the surface is simply connected, $c_0 = 0$ is necessary to avoid the singularity at $r = 0$. Then using Eq. (16) the diagonal elements of the surface tension tensor along meridian and azimuthal directions are obtained as

$$\sigma_1^1 = \frac{\Delta P}{2\kappa_2^2}, \tag{19}$$

$$\sigma_2^2 = \frac{\Delta P}{\kappa_2^2}\left(1 - \frac{\kappa_1^1}{2\kappa_2^2}\right). \tag{20}$$

Furthermore, solving Eq. (15), the shear component of the in-plane tension is obtained as

$$\sigma_1^2 = -\frac{c_1}{r^3}, \tag{21}$$

where $c_1$ is a constant of integration. For simply connected axisymmetric surfaces, $c_1 = 0$ is necessary due to the singularity at $r = 0$. As $\sigma_2^1 = \sigma_1^2 = 0$, $\sigma_1^1$ and $\sigma_2^2$ are then the principal stresses along meridian and azimuthal directions. Thus, for simply connected surfaces of revolution, the membrane balance equations can be solved directly just from the deformed shape and pressure. The edge tension $\boldsymbol{t}$ is not required in this case as the integration constants can be eliminated based on the singularity at $r = 0$. In fact, for simply connected axisymmetric shapes, $\sigma_1^1$ can be inferred directly at any location by using balance of forces acting on the disc normal to the rotation axis at that point. For the special case of a spherical membrane of radius $R$, $\sigma = \sigma_1^1 = \sigma_2^2$ and $\kappa_1^2 = \kappa_2^2 = 1/R$, which results in the Young-Laplace relation $\sigma = \Delta P R/2$.

##### Inverse elastostatics approach

The static determinacy of the balance equations can also be interpreted as invariance of membrane stress with respect to the constitutive behavior of the membrane. This constitutive invariance can be leveraged to setup an inverse elastostatics problem to solve for $\boldsymbol{\sigma}$. In this approach, an arbitrary constitutive behavior is assumed for the membrane material and starting from the known deformed configuration (including the external forces $\Delta P$ and $\boldsymbol{t}$ on $\partial\Gamma$) the stress-free reference configuration is treated as an unknown. The stresses in deformed configurations are determined by post-processing the results using the reference and deformed configurations and the assumed constitutive relation. Since in principle the stress inference problem is statically determinate, the estimated stresses should be independent of the chosen material model.

However, in practical applications, such as *in vivo* surface stress measurement of brain aneurysms [12, 13], $\boldsymbol{t}$ is often not known. For such cases, rather than Neumann boundary conditions, Dirichlet boundary conditions can be imposed on $\partial\Gamma$ to solve the inverse elastostatics problem. With Dirichlet boundary conditions, numerical experiments show that the effect of the choice of constitutive relation is weak and limited to the vicinity of the fixed edge, while invariance with respect to the constitutive relation is accurately satisfied far from the boundary [12, 14]. Thus, the stresses inferred by the inverse elastostatics approach with fixed boundary weakly depend on the choice of the constitutive behavior [15].

**Forward elastostatics penalty approach**

Another way of exploiting the constitutive invariance is proposed in Ref. [15] based on a penalty-like approach. The deformed membrane shape itself is taken as the stress-free reference configuration and the membrane is assumed to be sufficiently stiff such that after applying the inflation pressure the deformed shape remains close to the target shape. The advantage of this approach is the ease of implementation in a standard (forward) finite elements code. However, the choice of material parameters can be delicate. Moreover, Dirichlet boundary conditions imposed at the free edge are expected to perturb the solution close to the fixed edge in a way that depends on the constitutive parameters.

Both inverse and forward elastostatic approaches have been applied to the estimation of membrane stresses in biomedical applications and also to probe the point-wise constitutive response from inflation tests of soft tissues. However, obtaining the solution is contingent upon the existence of a stress-free reference configuration for the given choice of material model, parameters, and edge boundary conditions. Hence, the application of these methods to epithelial sheets may require using a more pertinent constitutive model for such active materials with an active pre-tension. Therefore, a direct approach not invoking a constitutive relation is more attractive for estimating surface stresses on inflated epithelial domes.

**Direct approach**

Romo et al.[16] proposed a direct approach to estimate $\boldsymbol{\sigma}$ that does not assume any constitutive behavior for the membrane material. The three components of the symmetric stress tensor are treated as the unknowns of the problem stated using Eqs. (12) and (13). The governing equations are discretized on a surface triangulation such that force balance is established for every element in terms of the unknown stress components defined at nodes. Additional constraints can be imposed on the balance equations if the edge tension are partially or fully known on the boundary $\partial\Gamma$. This results in an overdetermined system of linear equations, as there are three equations per element and three unknowns per node and for a triangulation the number of elements is higher than the number of nodes. Thus, a solution for $\boldsymbol{\sigma}$ can be obtained in a least-squares sense even if the original problem is ill-posed. We note that restricting the degrees of freedom to be less than the equations introduces an implicit regularization and provides a solution that minimizes the least-squares error. The discretization scheme proposed in Ref. [16] was motivated by force balance at the centroid of every triangular element.

#### 2.2 Inverse problem formulation of cMSM

Here, we present a systematic formulation of the direct method discussed above and its regularization.

The balance equations (12) and (13) can be combined and re-expressed in a convenient vectorial form as [8, 9]

$$\frac{1}{\sqrt{g}}(\sqrt{g}\sigma^{ab}\boldsymbol{e}_a)_{,b} + \Delta P\boldsymbol{n} = 0, \tag{22}$$

where $g = \det g_{ab}$. Note that the unknown in our inverse problem is the symmetric surface tensor $\boldsymbol{\sigma}$ while $\Delta P$ and the deformed surface $\Gamma$ is known *i.e.* $\boldsymbol{e}_1$, $\boldsymbol{e}_2$, and $\boldsymbol{n}$ are given. We intend to use finite elements to obtain the discretized form of the governing equations. To this end, we can obtain the following weak form of the balance equations using an arbitrary vector test function $\boldsymbol{w}$

$$\int_\Gamma \left[\frac{1}{\sqrt{g}}(\sqrt{g}\sigma^{ab}\boldsymbol{e}_a)_{,b} + \Delta P\boldsymbol{n}\right]\cdot\boldsymbol{w}\,dS = 0. \tag{23}$$

To obtain the discrete form of Eq. (23), the surface $\Gamma$ is discretized into finite elemental domains by triangulation. For inflated dome shapes that are of interest in this study, we can obtain a global parametrization for the open surface $\Gamma$ with the basal footprint edge $\partial\Gamma$ using well-established parametrization methods for triangulated surfaces such as disc conformal maps [17]. This global parametrization can then be used to calculate $\{\boldsymbol{e}_1, \boldsymbol{e}_2, \boldsymbol{n}\}$ at all nodes of the mesh and directly discretize the components of the surface tensor $\boldsymbol{\sigma}$ defined at nodes. However, in a general case where such a global parametrization is not available or not possible (closed surfaces), the surface geometry can be evaluated using the local elemental parametrization and techniques to discretize surface tensor fields [18].

Using the global parametrization of $\Gamma$, the components of surface tension tensor $\boldsymbol{\sigma}$ are expressed in the basis $\{\boldsymbol{e}_1, \boldsymbol{e}_2\}$ and can be interpolated using continuous basis functions, which may have discontinuous derivatives. However, given the form of Eq. (23), it is convenient instead to interpolate the entire term $\sqrt{g}\sigma^{ab}\boldsymbol{e}_a$ by defining two vectors $\boldsymbol{s}^1 = \sqrt{g}(\sigma^{11}\boldsymbol{e}_1 + \sigma^{21}\boldsymbol{e}_2)$ and $\boldsymbol{s}^2 = \sqrt{g}(\sigma^{12}\boldsymbol{e}_1 + \sigma^{22}\boldsymbol{e}_2)$ at all nodes. First, the global parametrization is used to define $\boldsymbol{e}_{I1}$, $\boldsymbol{e}_{I2}$, $\boldsymbol{n}_I$, and $g_I$ at a node $I$ using Eqns. (9), (11), and (10). The surface tension components at node $I$ are denoted as $\{\sigma_I^{11}, \sigma_I^{22}, \sigma_I^{12}\}$. Then the terms in Eq. (23) are interpolated as

$$\boldsymbol{s}^b = \sqrt{g}\sigma^{ab}\boldsymbol{e}_a(\boldsymbol{x}) = \sum_{J\in E} \boldsymbol{s}_J^b N_J \circ \boldsymbol{\psi}^{-1}(\boldsymbol{x}). \tag{24}$$

and

$$\boldsymbol{n}(\boldsymbol{x}) = \sum_{J\in E} \boldsymbol{n}_J N_J \circ \boldsymbol{\psi}^{-1}(\boldsymbol{x}) \tag{25}$$

where $\boldsymbol{x} = \psi(\boldsymbol{\chi})$ is the isoparametric mapping for the triangular element and $N_J(\boldsymbol{\chi})$ are the linear shape functions for node $J$ of element $E$ [19]. The integration over $\Gamma_e$ is performed numerically using Gaussian quadrature using isoparametric mapping of the triangular elements. The first term in Eq. (23) is evaluated as

$$(\sqrt{g}\sigma^{ab}\boldsymbol{e}_a)_{,b}(\boldsymbol{x}) = \sum_{J\in E} \boldsymbol{s}_J^b \frac{\partial N_J}{\partial \boldsymbol{\chi}} \frac{\partial \boldsymbol{\chi}}{\partial \xi^b} \circ \boldsymbol{\psi}^{-1}(\boldsymbol{x}). \tag{26}$$

Following Galerkin method, the test function can be expressed as

$$\boldsymbol{w}(\boldsymbol{x}) = \sum_{I\in E} \boldsymbol{w}_I N_I \circ \boldsymbol{\psi}^{-1}(\boldsymbol{x}), \tag{27}$$

where $\boldsymbol{w}_I$ are the nodal values of the test function. Using Eqns. (23), (24), (25), (26), and (27) along with arbitrariness of $\boldsymbol{w}_I$, we obtain three equations for every free node. Since we also have three unknowns per free node, this results in a square linear system of equations. The nodal array of unknowns is formed by combining the unknown tension components, $\mathsf{u}_J = [\sigma_J^{11}, \sigma_J^{22}, \sigma_J^{12}]$. The nodal unknowns are collected together in an array $\mathsf{u} = [\mathsf{u}_1, \mathsf{u}_2, ..., \mathsf{u}_N]^T$, where $N$ is the total number of nodes. The system of linear equations obtained from Eqs. (23) are expressed as $\mathsf{Au} = \mathsf{b}$, $\mathsf{A}$ is a square matrix of size $3N \times 3N$ and $\mathsf{b}$ is a column vector of size $3N \times 1$ resulting from the pressure term in Eq. (23).

##### 2.2.1 Regularization

The inverse problem of force inference from the deformation field is often ill-posed. For instance, in traction force microscopy (TFM) [20], the traction forces at the cell-matrix interface are inferred from displacement of fidicuary markers embedded in the soft matrix. The inversion process is inherently unstable due to the slow decay of the elasticity Green's function *i.e.* noise in the measured displacements results in large changes in the inferred traction field. The inverse problem is therefore often regularized by minimizing the force balance equations along with a zeroth-order Tikhonov regularization that penalizes the $L_2$-norm of the unknown forces. The regularization term is scaled by a scalar regularization parameter $\lambda$ which is chosen so as to establish a balance between regularizing the solution and satisfying the force balance equations. The parameter $\lambda$ can be chosen based on the L-curve method [20], which can be subjective as an obvious inflection point may not always be present, or more objectively based on Bayesian inference [21]. Apart from $L_2$-regularization, other ways of penalizing undesirable features of traction fields have also been explored for TFM including $L_1$-regularization and gradient-based penalties [21].

The inverse problem of membrane tension inference is different than TFM in that we are not solving an ill-posed inverse elasticity problem. In TFM, the in-plane momentum balance equations alone are not sufficient to infer tractions and constitutive equations for the soft matrix are necessary to formulate the inverse problem. In the membrane tension inference problem, the out-of-plane balance relation in Eq. (13) renders the system statically determinate and allows one to directly solve for surface stresses. Yet the inverse problem can become ill-posed if the given surface shape cannot support a pressure through a membrane stress, e.g. if it is locally planar, reflecting the sensitivity of the solution to shape and its variations inherent to the shape acquisition method based on confocal stacks. Indeed, in practice we observe that $\mathsf{Au} = \mathsf{b}$ is ill-conditioned even for simple axisymmetric shapes such as a spherical cap (Supplementary Fig. 4). Regularization is therefore necessary to solve this inverse problem. It amounts to minimizing the following function with respect to $\mathsf{u}$,

$$E(\mathsf{u}) = \frac{1}{2}||\mathsf{Au} - \mathsf{b}||^2 + L(\mathsf{u}), \tag{28}$$

where $L(\mathsf{u})$ is the discretized regularization contribution that penalizes any undesirable characteristics in the unknown tension field. A natural first choice for regularization is a zeroth-order Tikhonov regularization term that penalizes magnitude of surface tensions expressed as

$$L_0(\boldsymbol{\sigma}) = \frac{\lambda_0^2}{2}\int_\Gamma \boldsymbol{\sigma} : \boldsymbol{\sigma}\, dS, \tag{29}$$

where $\lambda_0$ is a dimensionless regularization parameter. However, surface stresses inferred using only this regularization were observed to exhibit sharp gradients in surface tensions. We therefore did not include this regularization term in our analysis.

We then considered a first-order regularization of the form

$$L_{1t}(\boldsymbol{\sigma}) = \frac{\lambda_t^2}{2}\int_\Gamma \boldsymbol{\nabla}\text{trace}(\boldsymbol{\sigma}) \cdot \boldsymbol{\nabla}\text{trace}(\boldsymbol{\sigma})\, dS = \frac{\lambda_t^2}{2}\int_\Gamma \sigma^a_{a|e}\sigma^b_{b|f} g^{ef}\, dS, \tag{30}$$

where the regularization parameter $\lambda_t$ provides a length scale to penalize tension gradients. Here $g^{ef}$ are the components of the inverse of the metric tensor and $\sigma^b_{b|e}$ denotes the components of covariant derivative of trace($\boldsymbol{\sigma}$) [7, 9]. This regularization penalizes gradients in the mean surface tension.

We further found that penalizing only $\boldsymbol{\nabla}\mathrm{trace}(\boldsymbol{\sigma})$ does not necessarily restrain sharp rotations (swirl) in the surface tension field. To regularize such features the following term penalizing the curl of surface tension is introduced

$$L_{1c}(\boldsymbol{\sigma}) = \frac{\lambda_c^2}{2}\int_\Gamma \mathrm{curl}\,\boldsymbol{\sigma} : \mathrm{curl}\,\boldsymbol{\sigma}\, dS = \frac{\lambda_c^2}{2}\int_\Gamma \epsilon^{ge}\sigma^{ab}_{|e}\epsilon^{hf}\sigma^{cd}_{|f}\, g_{ac}\, g_{bd}\, g_{gh}\, dS, \tag{31}$$

where $\boldsymbol{\epsilon}$ is the Levi-Civita tensor [7].

The first-order regularization was found to work best for all cases analyzed in this study. The discrete version of the regularization terms $L_{1t}$ and $L_{1c}$ are obtained by using linear elements to interpolate nodal values of surface tension $\sigma_I^{ab}$ and covariant basis vectors $\boldsymbol{e}_{I1}$ and $\boldsymbol{e}_{I2}$:

$$L(\mathsf{u}) = \frac{1}{2}\mathsf{u}^T\left(\lambda_t^2\mathsf{Q}_t + \lambda_c^2\mathsf{Q}_c\right)\mathsf{u} = \frac{\lambda_t^2}{2}\mathsf{u}^T\mathsf{Q}\mathsf{u}, \tag{32}$$

where $\mathsf{Q}_t$ and $\mathsf{Q}_c$ are the regularization matrices resulting from discretizing $L_{1t}(\boldsymbol{\sigma})$ and $L_{1c}(\boldsymbol{\sigma})$, and

$$\mathsf{Q} = \mathsf{Q}_t + \frac{\lambda_c^2}{\lambda_t^2}\mathsf{Q}_c.$$

The surfaces fit to the experimental point clouds may have regions that are flat (zero Gaussian curvature) or concave (negative Gaussian curvature). Such regions exhibit negative principal tensions that are not compatible with a stable membrane state of stress. The regularization terms considered above do not restrict negative principal tensions. Thus when analyzing experimental data we additionally impose the following inequality constraints

$$\det\boldsymbol{\sigma}_J = \sigma_J^{11}\sigma_J^{22} - \left(\sigma_J^{12}\right)^2 > 0$$

for each node $J$ when minimizing Eq. (28). The minimization problem however becomes nonlinear after introducing the inequality constraint and is solved using the constrained minimization function (`fmincon`) in Matlab.

Taken together, application this inverse approach to curved epithelia under luminal pressure is termed as curved Monolayer Stress Microscopy (cMSM), which can be viewed as a generalization of the planar MSM [22, 23]. However, in cMSM, any assumptions about the constitutive behavior of the membrane are not necessary.

#### 2.3 Testing the cMSM inverse approach

##### Axisymmetric shapes

We systematically verified our approach by reconstructing surface tensions on axisymmetric shapes that can be directly compared with the closed-form solution given by Eqs. (19) and (20). The difference between the inferred tension $\boldsymbol{\sigma}$ and the closed-form solution $\boldsymbol{\sigma}_{\mathrm{CF}}$ at all nodes is quantified using the relative error evaluated as $||\boldsymbol{\sigma} - \boldsymbol{\sigma}_{\mathrm{CF}}||/||\boldsymbol{\sigma}_{\mathrm{CF}}||$.

We analyzed ellipsoidal caps with the aspect ratio given as $\alpha = r_a/r_b$, where $r_b$ and $r_a$ are the principal radii. For each case, the inverse problem is solved for a wide range of values of $\lambda_t$ which regularizes the gradients in tension. Regularization in the curl of surface tension is controlled by the parameter $\lambda_c$. This analysis provides guidance to choose the regularization parameters when analyzing the experimentally obtained membrane shapes. For all cases, the circular footprint is set to have radius $r_b = 40\,\mu\mathrm{m}$ to be comparable to the experimental domes, while $\Delta P = 25\,\mathrm{Pa}$ such that $\mathrm{trace}(\boldsymbol{\sigma}) = 1\,\mathrm{mN/m}$ for a spherical dome.

Surface tension recovery for a spherical cap ($\alpha = 1$) is shown in Supplementary Fig. 5. In this and other figures, the symmetric stress tensor is represented graphically in terms of its mutually orthogonal principal directions of stress (eigenvectors) and principal stresses along these directions (eigenvalues), represented by orthogonal pairs of arrows whose length is proportional to the magnitude of the corresponding principal tensions and whose divergence/convergence represent their positive/negative sign. The color of the surface shows the hydrostatic component of the surface tension *i.e.* trace($\boldsymbol{\sigma}$). For a spherical cap in Supplementary Fig. 5, an isotropic state of stress is represented by two mutually orthogonal pairs of arrows of the same length.

The inverse problem is solved for a fixed $\lambda_c$ and a range of $\lambda_t$ values acting as a regularization parameter. The tension obtained at every node is compared with the expected analytical solution to quantify the error as shown in Supplementary Fig. 5a. With $\lambda_c = 0$, the relative error is within 1% for $\lambda_t$ roughly between 0.002 and 0.1. For $\lambda_t$ below this range the problem is under-regularized, which leads to larger errors, whereas for $\lambda_t$ above this range the problem is over-regularized, which again leads to larger errors. Instead of performing a component-wise comparison of $\boldsymbol{\sigma}$ with the analytical solution, if only trace($\boldsymbol{\sigma}$) is compared the errors are much smaller over a wider range of regularization parameter values (Supplementary Fig. 6). This is expected since $\lambda_t$ explicitly penalizes gradients in trace($\boldsymbol{\sigma}$).

Introducing the curl-based regularization with $\lambda_c$ is observed to further improve the overall recovery of surface tensions (Supplementary Fig. 5a). For each $\lambda_c$, the $\lambda_t$ value corresponding to the minimum error in Supplementary Fig. 5a can be clearly associated with the corner in the L-curve, where the regularization term is plotted against the residual forces (Supplementary Fig. 5d). The regularization term is evaluate as $\sqrt{\mathsf{u}^T\mathsf{Q}\mathsf{u}/A}$, where $A$ is the total dome area. The residual forces are quantified using $||\mathsf{A}\mathsf{u} - \mathsf{b}||/||\mathsf{b}||$. This corner point can also be identified as the regularization parameter value $\lambda_t$ after which the residual starts increasing as shown in Supplementary Fig. 5c.

Note that for $\lambda_t < \lambda_c$, the regularization is dominated by the curl-based term, which improves the solutions obtained for low $\lambda_t$ values. In general, we observe that introducing $\lambda_c$ penalizes short wavelength features in $\boldsymbol{\sigma}$, which are not necessarily filtered through $\lambda_t$ regularization. Moreover, choosing $\lambda_t$ for a given $\lambda_c$ from Figs. 5c or 5d is less ambiguous compared to when $\lambda_c = 0$.

A spherical cap is a special case as the curvature is isotropic and constant, leading to constant and isotropic surface tensions everywhere. Thus, any $\lambda_c > 0$ can be observed to significantly improve the solution (Supplementary Fig. 5a). However, for shapes with significant curvature gradients, we expect tension gradients as well. Using a higher value of $\lambda_c$ in such a case might overly-penalize the tension gradients leading to larger errors. Indeed, this becomes apparent when the surface stress recovery is performed on prolate and oblate ellipsoidal caps as shown in Figs. 7 and 8. For instance, $\lambda_c = 0.01$ seems to work well for oblate ellipsoids, while increasing $\lambda_c$ to 0.1 leads to slightly larger errors.

##### Inflated hyperelastic membrane

To further test the approach beyond axisymmetric shapes, we generated deformed shapes by inflating a flat patch of a neo-Hookean membrane with fixed edges. The basal footprints were chosen to be elliptical with aspect ratios 1, 2, and 3. The smaller principal radius of the elliptical footprints is fixed at 25 µm. The strain energy per unit reference area for the neo-Hookean membrane is of the form

$$\psi(I, J) = \mu(I - 2) + \lambda(J - 1)^2,$$

where $I = \mathrm{trace}(\boldsymbol{C})$, $J = \sqrt{\det \boldsymbol{C}}$, and $\boldsymbol{C}$ is the right Cauchy-Green deformation tensor [9]. The material parameters are $\mu = 1\,\mathrm{mN/m}$ and $\lambda = 2\mu$, while pressure applied is $\Delta P = 400\,\mathrm{Pa}$.

The inflation problem was solved using finite elements and membrane stresses were computed at mesh nodes by post-processing the solution. The inferred membrane stresses obtained by solving the inverse problem on the given deformed shapes are then compared with the finite element solution as shown in Figs. 9, 10, and 11. In all cases, the surface tensions are recovered to be within $\sim 3\%$ of the known numerical solution.

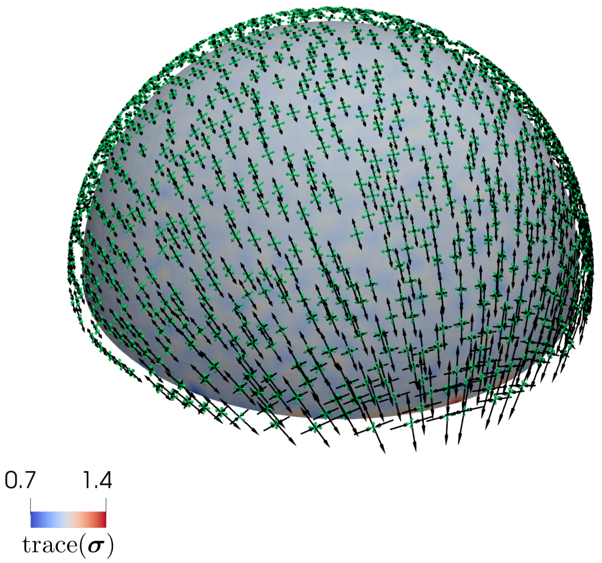

Supplementary Figure 4: Membrane stress inference on a spherical cap without any regularization. The black arrows represent the inferred membrane stresses and green arrows represent the expected solution for a spherical cap.

In each example, for a given value of $\lambda_c$ we sweep for a wide range of $\lambda_t$. The choice of $\lambda_t$ is then based on the corner where the residual starts sharply increasing with $\lambda_t$ (Supplementary Fig. 9c). Note that this corner point also corresponds to the minimum relative error shown in Supplementary Fig. 9a and the point where the residual starts increasing with $\lambda_t$ in Supplementary Fig. 9b. When analyzing experimental data for which the true solution is not known, this approach based on a plot similar to Figs. 9b and c is used to select the regularization parameter.

#### 2.4 Surface stress inference on epithelial domes

##### Dome segmentation

To extract the dome shape from experimental data, we generate a three-dimensional point cloud corresponding to the basal cell faces (Supplementary Fig. 12). First, the $x$–$y$ slice located roughly 2–4 µm above the substrate is identified and segmented as the dome footprint. Using a $x$–$y$ slice very close to the substrate can include protrusions extended by substrate bound cells that may distort the basal footprint shape. For a given slice, points are manually selected that denote the basal cell faces. A spline fit is obtained to the selected points which is used to generate the final point cloud with uniformly spaced points. The $x$–$z$ and $y$–$z$ slices spaced roughly 7.5 µm apart are then processed in a similar manner. The point cloud for a given dome is obtained by combining the points from all segmented slices.

The steps involved in fitting a surface $\Gamma$ to the point cloud obtained from experimental data are shown in Supplementary Fig. 13. The three-dimensional point cloud (Supplementary Fig. 12b) is first triangulated (Supplementary Fig. 13a). Note that the point cloud shown in Supplementary Fig. 12b is down-sampled to aid the triangulation process. Then, the open surface is mapped on a unit disc using disc a conformal map [17], which provides a global parametrization for $\Gamma$ (Supplementary Fig. 13b). The part of the unit disc corresponding to the inflated dome surface is re-meshed using triangular elements to improve the mesh quality (Supplementary Fig. 13c).

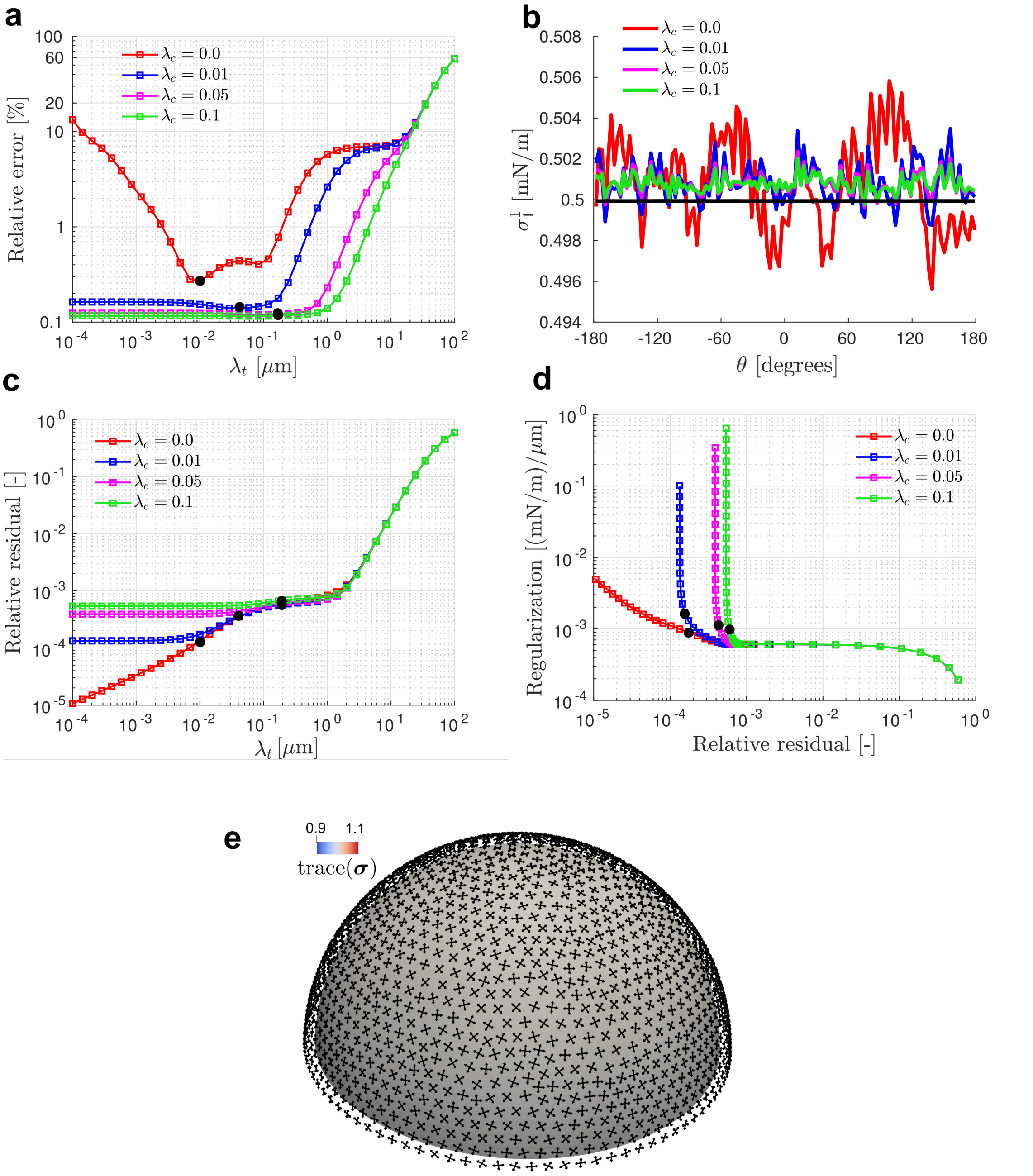

Supplementary Figure 5: Surface tension inference on a spherical cap. The inverse problem solution is analyzed as a function of the regularization parameter $\lambda_t$ for $\lambda_c = \{0.0, 0.01, 0.05, 0.1\}$. (a) The relative error between the inferred tension and the closed-form solution is sufficiently low ($< 1\%$) for a wide range of $\lambda_t$ values which depends on the choice of $\lambda_c$. (b) The tangential component of the surface tension along the edge $\sigma_1^1$ is shown as a function of angle of revolution $\theta$. The black line is the closed-form solution. (c) The residual of the nodal force balance equations as a function of $\lambda_t$ clearly shows that the optimal regularization parameter is associated with the corner point where residual starts increasing. (d) Regularization functional plotted against the residual, in the so-called L-curve. The corner in the L-curve can be identified with the optimal regularization parameter $\lambda_t$ for a given $\lambda_c$. (e) Surface stress solution obtained for $\lambda_c = 0.1$.

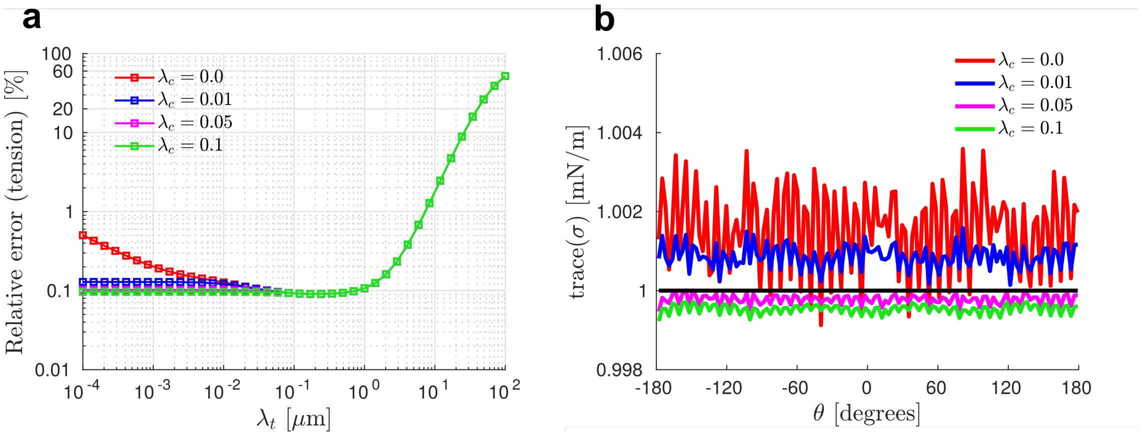

Supplementary Figure 6: (a) Relative error in the inferred tension trace($\boldsymbol{\sigma}$) and closed-form solution trace($\boldsymbol{\sigma}_{\mathrm{CF}}$) for spherical cap shown in Supplementary Fig. 5. (b) The tension on the edge is compared with the close-form solution shown as a solid black line.

This triangulation is then fitted back to the point cloud shown in Supplementary Fig. 13a with a smoothness penalty [24] to obtain a smooth surface fit (Supplementary Fig. 13d).

##### Regularization parameter selection

For all experimental data analyzed in this study, we use $\lambda_c = 0.1$ based on the examples presented above, specifically the hyperelastic membrane inflation simulations. A somewhat higher regularization is expected and intentional when working with experimental data due to measurement and membrane-approximation errors, especially at low inflation levels. A representative example of analysis performed on an experimentally obtained surface is shown in Supplementary Fig. 14. The regularization parameter $\lambda_t$ is chosen as shown in Supplementary Fig. 14 following the same procedure outlined in the previous section.

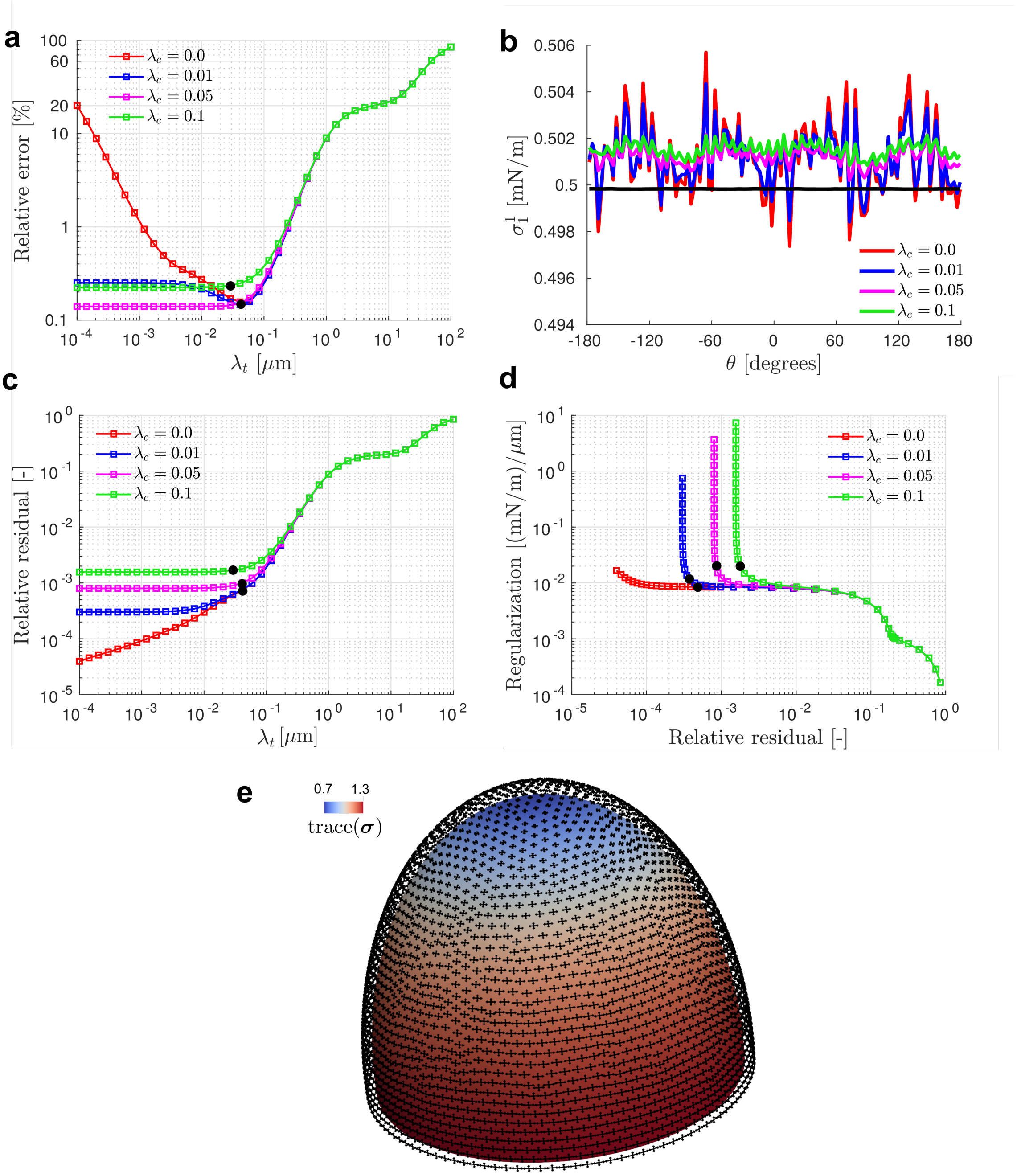

Supplementary Figure 7: Surface tension inference on a prolate cap with aspect ratio $\alpha = 1.5$. The inverse problem solution is analyzed as a function of the regularization parameter $\lambda_t$ for $\lambda_c = \{0.0, 0.01, 0.05, 0.1\}$. (a) The relative error between the inferred tension and the closed-form solution is sufficiently low ($< 1\%$) for a wide range of $\lambda_t$ values. (b) The tangential component of the surface tension along the edge $\sigma_1^1$ is shown as a function of angle of revolution $\theta$. The black line represents the closed-form solution. (c) The residual of the nodal force balance equations as a function of $\lambda_t$ clearly shows that the optimal regularization parameter is associated with the corner point where residual starts increasing. (d) The corner in the L-curve can be identified with the optimal regularization parameter $\lambda_t$ for a given $\lambda_c$. (e) Surface stress solution obtained for $\lambda_c = 0.05$.

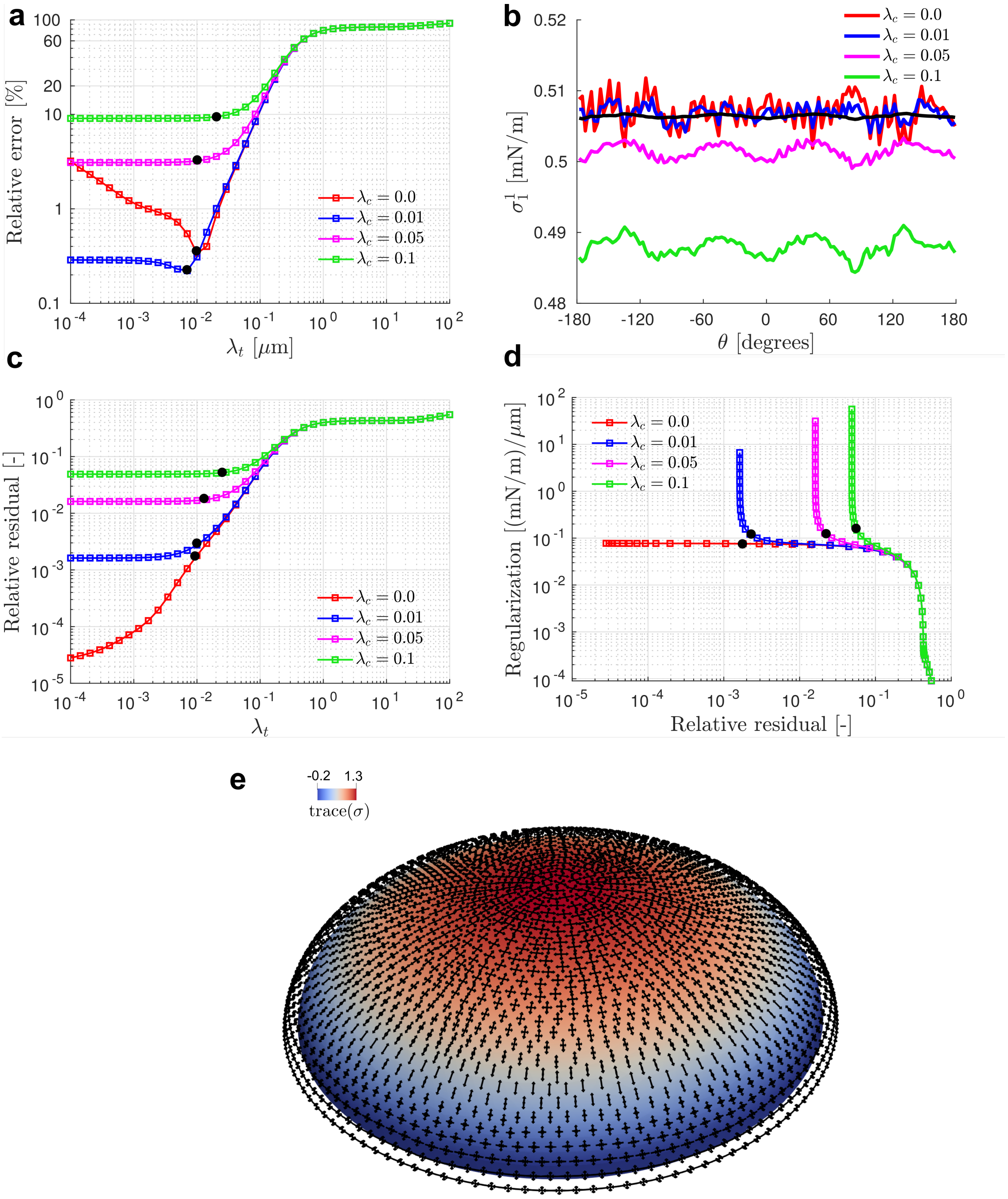

Supplementary Figure 8: Surface tension inference on an oblate cap with aspect ratio $\alpha = 0.5$. The inverse problem solution is analyzed as a function of the regularization parameter $\lambda_t$ for $\lambda_c = \{0.0, 0.01, 0.05, 0.1\}$. (a) The relative error between the inferred tension and the closed-form solution is sufficiently low ($< 1\%$) for a much smaller range of $\lambda_t$ values compared to Figs. 5 and 7. For $\lambda_c = 0.1$ the problem is over-regularized which results in higher errors. (b) The tangential component of the surface tension along the edge $\sigma_1^1$ is shown as a function of angle of revolution $\theta$. The black line represents the closed-form solution. (c) The residual of the nodal force balance equations as a function of $\lambda_t$ clearly shows that the optimal regularization parameter is associated with the corner point where residual starts increasing. (d) The corner in the L-curve can be identified with the optimal regularization parameter $\lambda_t$ for a given $\lambda_c$. (e) Surface stress solution obtained for $\lambda_c = 0.01$.

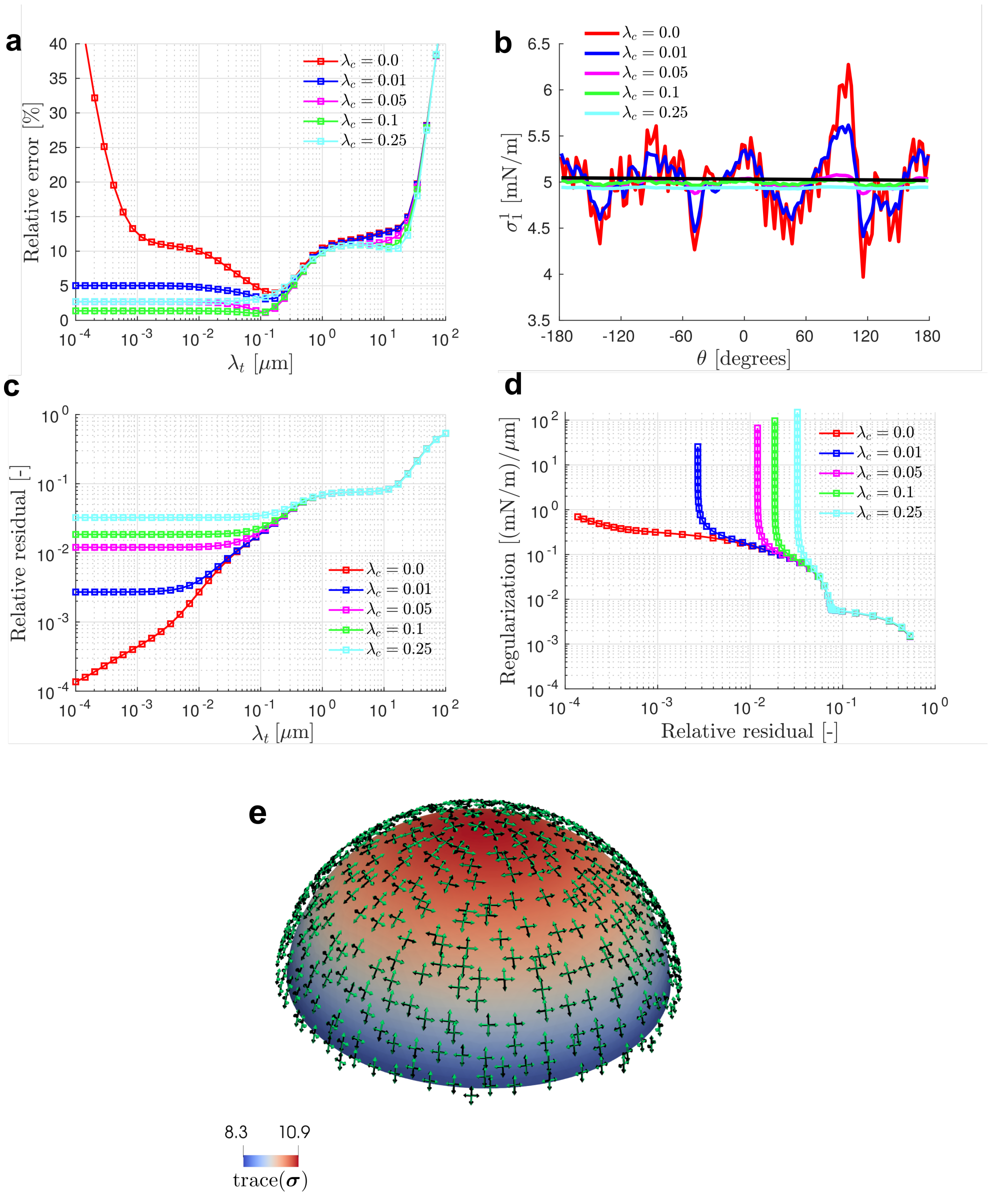

Supplementary Figure 9: Surface tension inference on an inflated Neohookean membrane with a circular footprint. The inverse problem solution is analyzed as a function of the regularization parameter $\lambda_t$ for $\lambda_c = \{0.0, 0.01, 0.05, 0.1, 0.25\}$. (a) The relative error between the inferred tension and the finite element solution is minimized for $\lambda_t \approx 0.1$. (b) The tangential component of the surface tension along the edge $\sigma_1^1$ is shown as a function of angle of revolution $\theta$. The solution is plotted for $\lambda_t$ corresponding to the minimum error in (a). The black line is the finite element solution. (c) The residual of the nodal force balance equations as a function of $\lambda_t$ clearly shows that the optimal regularization parameter is associated with the corner point where residual starts increasing. (d) The corner in the L-curve can also be identified with the optimal regularization parameter $\lambda_t$ for a given $\lambda_c$. (e) Surface stress solution obtained for $\lambda_c = 0.1$.

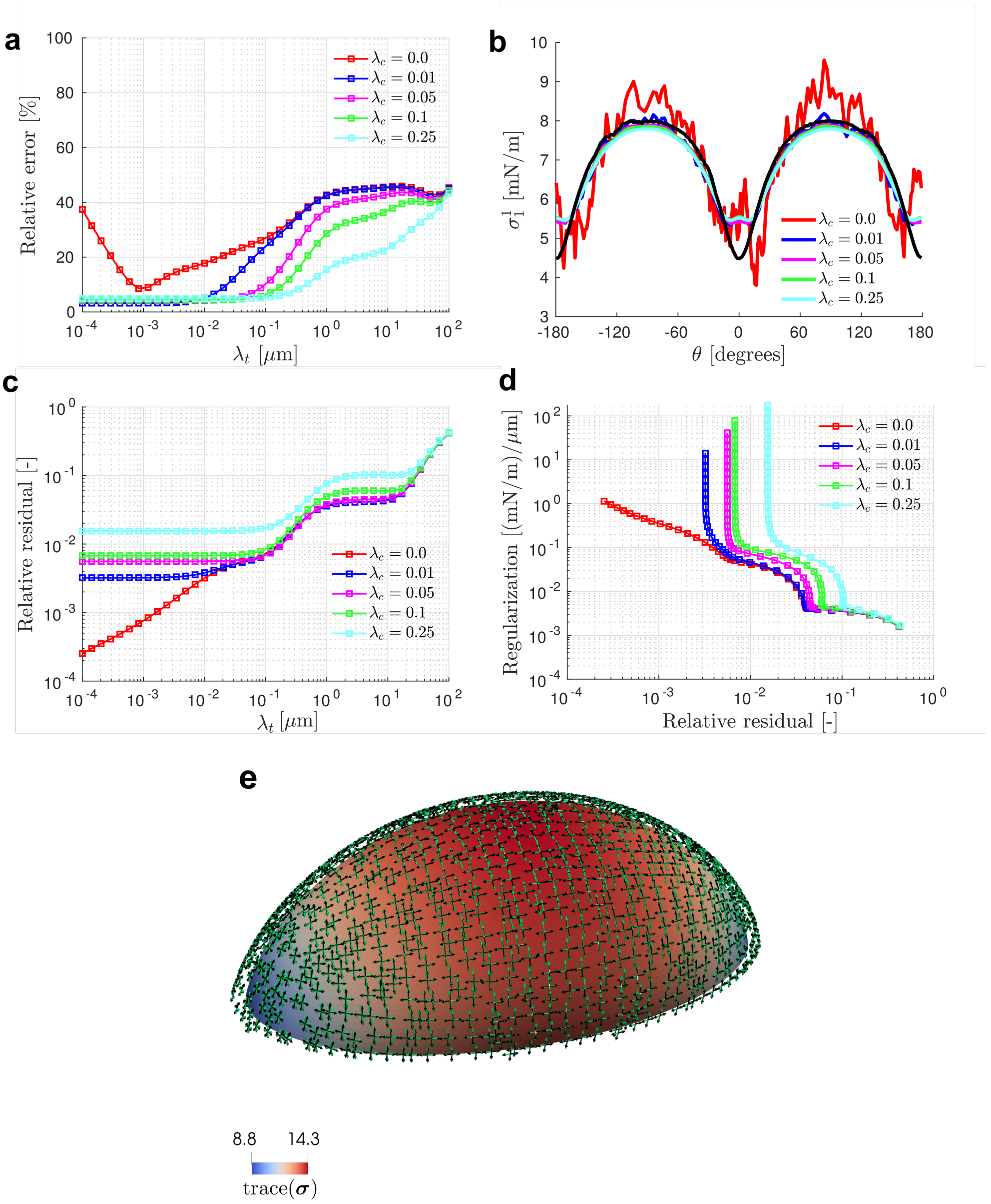

Supplementary Figure 10: Surface tension inference on a an inflated neohookean membrane with an elliptical footprint of aspect ratio 2. The inverse problem solution is analyzed as a function of the regularization parameter $\lambda_t$ for $\lambda_c = \{0.0, 0.01, 0.05, 0.1, 0.25\}$. (a) The relative error between the inferred tension and the finite element solution is shown as a function of $\lambda_t$. (b) The tangential component of the surface tension along the edge $\sigma_1^1$ is shown as a function of angle of revolution $\theta$. The solution is plotted for $\lambda_t$ corresponding to the minimum error in (a). The black line is the finite element solution. (c) The residual of the nodal force balance equations as a function of $\lambda_t$ clearly shows that the optimal regularization parameter is associated with the corner point where residual starts increasing. (d) The corner in the L-curve can also be identified with the optimal regularization parameter $\lambda_t$ for a given $\lambda_c$. (e) Surface stress solution obtained for $\lambda_c = 0.1$.

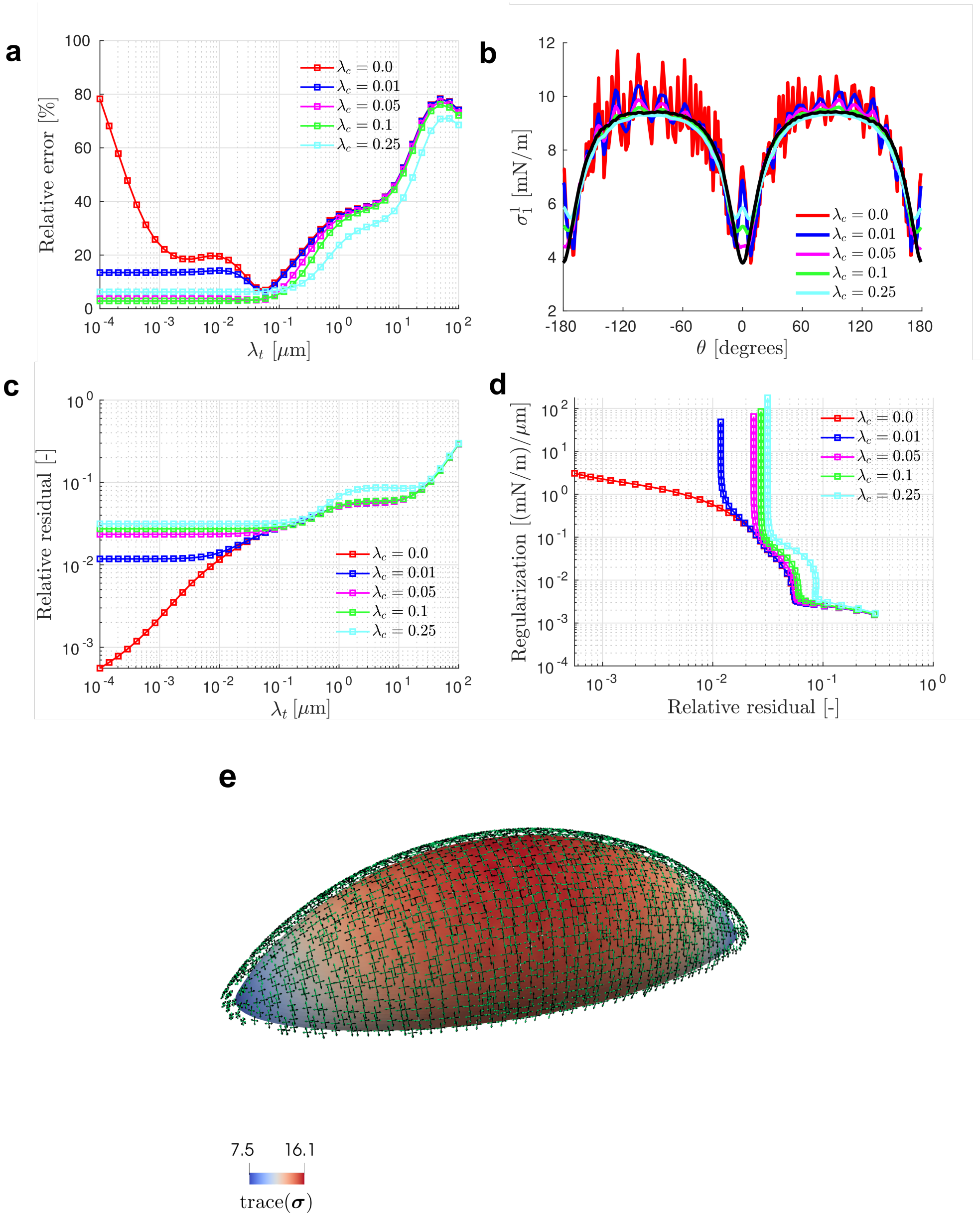

Supplementary Figure 11: Surface tension inference on a an inflated neohookean membrane with an elliptical footprint of aspect ratio 3. The inverse problem solution is analyzed as a function of the regularization parameter $\lambda_t$ for $\lambda_c = \{0.0, 0.01, 0.05, 0.1, 0.25\}$. (a) The relative error between the inferred tension and the finite element solution is shown as a function of $\lambda_t$. (b) The tangential component of the surface tension along the edge $\sigma_1^1$ is shown as a function of angle of revolution $\theta$. The solution is plotted for $\lambda_t$ corresponding to the minimum error in (a). The black line is the finite element solution. (c) The residual of the nodal force balance equations as a function of $\lambda_t$ clearly shows that the optimal regularization parameter is associated with the corner point where residual starts increasing. (d) The corner in the L-curve can also be identified with the optimal regularization parameter $\lambda_t$ for a given $\lambda_c$. (e) Surface stress solution obtained for $\lambda_c = 0.1$.

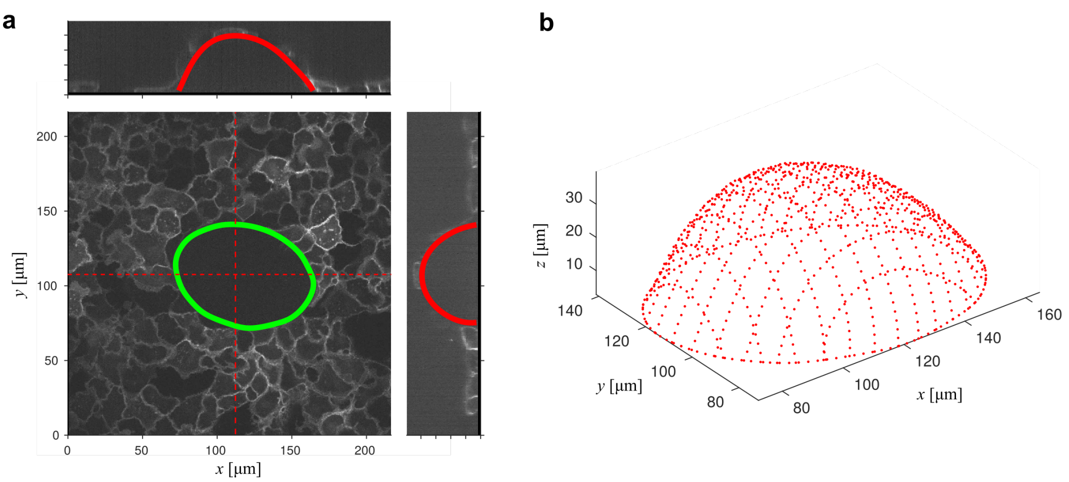

Supplementary Figure 12: (a) Segmentation of dome by selecting points on $x$–$y$, $x$–$z$, and $y$–$z$ slices of the image stack. Spline fits obtained to the selected points are shown for the basal footprint and selected $x$–$z$ and $y$–$z$ planes. (b) The point cloud representing the extracted luminal surface.

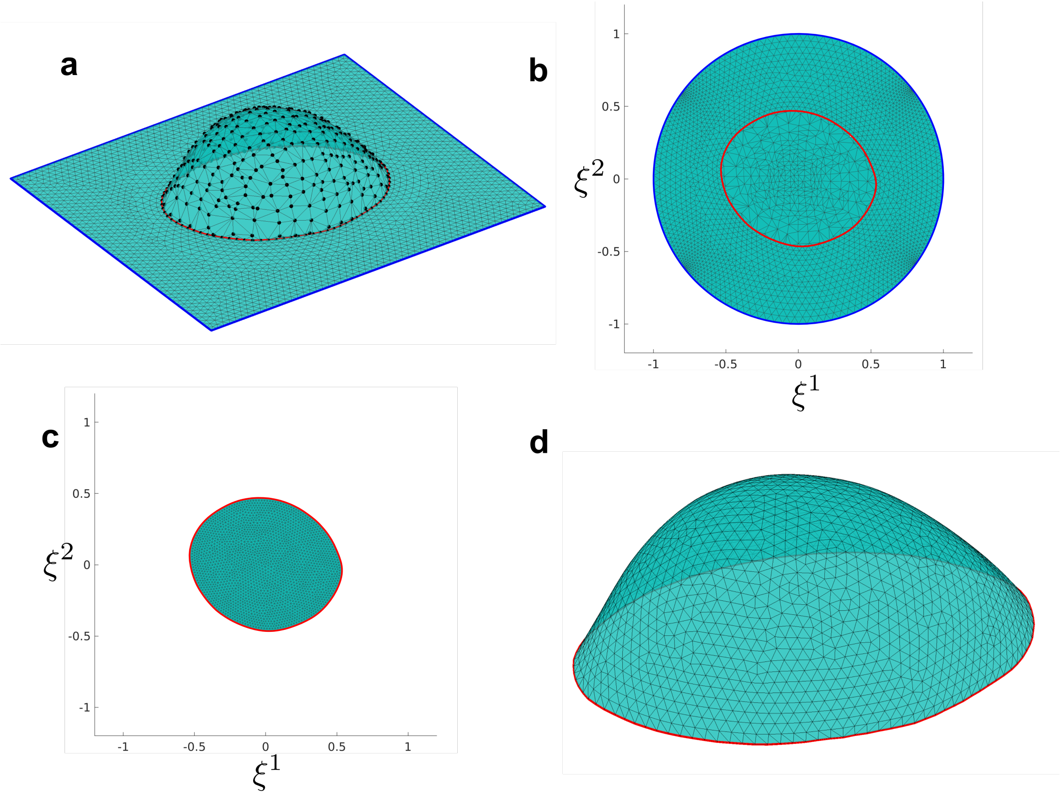

Supplementary Figure 13: Steps used to fit a smooth surface to the point cloud of epithelial dome basal lumen. (a) The point cloud (black points) is triangulated to generate the surface mesh for the inflated dome. The dome footprint is shown as the red line. (b) The surface triangulation in (a) is mapped on a unit disk using a disc conformal map. (c) Re-meshed dome surface in the parametric domain. (d) The mesh shown in (c) is fitted to the point cloud shown in (a) with a smoothness penalty.

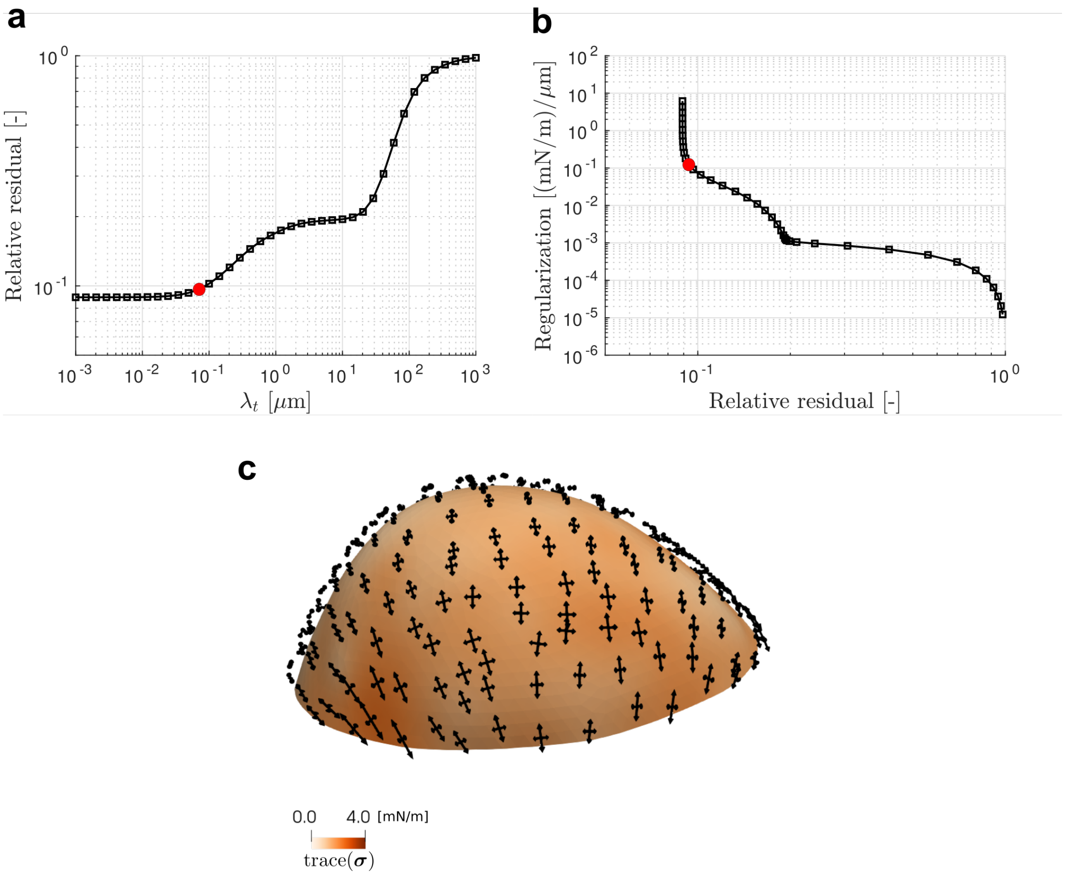

Supplementary Figure 14: Surface stress inference on epithelial dome surface obtained in Supplementary Fig. 13. The regularization parameter $\lambda_t$ is determined from (a), which can be identified as the corner point on the L-curve shown in (b). The chosen point is shown in both (a) and (b). (c) The inferred surface stresses for the chosen regularization parameters, $\lambda_c = 0.1$ and $\lambda_t = 0.07$.

### Supplementary references

1. Alt, S., Ganguly, P. & Salbreux, G. Vertex Models : from Cell Mechanics to Tissue Morphogenesis. *Philos. Trans. R. Soc. Lond. B Biol. Sci.* **372,** 20150520 (2017).
2. Belytschko, T., Liu, W. K., Moran, B. & Elkhodary, K. *Nonlinear Finite Elements for Continua and Structures* en (John Wiley & Sons, 2014).
3. Harris, A. R. *et al.* Characterizing the mechanics of cultured cell monolayers. *Proceedings of the National Academy of Sciences* **109,** 16449–16454 (2012).
4. Pérez-González, C. *et al.* Mechanical compartmentalization of the intestinal organoid enables crypt folding and collective cell migration. en. *Nat. Cell Biol.* **23,** 745–757 (2021).
5. Latorre, E. *et al.* Active superelasticity in three-dimensional epithelia of controlled shape. *Nature* **563,** 203–208 (2018).
6. Fouchard, J. *et al.* Curling of epithelial monolayers reveals coupling between active bending and tissue tension. en. *Proc. Natl. Acad. Sci. U. S. A.* **117,** 9377–9383 (2020).
7. Do Carmo, M. P. *Differential Geometry of Curves and Surfaces* en (Courier Dover Publications, 2016).
8. Naghdi, P. M. in *Linear Theories of Elasticity and Thermoelasticity: Linear and Nonlinear Theories of Rods, Plates, and Shells* (ed Truesdell, C) 425–640 (Springer Berlin Heidelberg, Berlin, Heidelberg, 1973).
9. Marsden, J. E. & Hughes, T. J. R. *Mathematical Foundations of Elasticity* en (Dover Publications, New York, New York, USA, 1994).
10. Hill, R. A theory of the plastic bulging of a metal diaphragm by lateral pressure. *The London, Edinburgh, and Dublin Philosophical Magazine and Journal of Science* **41,** 1133–1142 (1950).
11. Humphrey, J. D. & Kyriacou, S. K. The use of Laplace's equation in aneurysm mechanics. en. *Neurol. Res.* **18,** 204–208 (1996).
12. Lu, J., Zhou, X. & Raghavan, M. L. Inverse method of stress analysis for cerebral aneurysms. en. *Biomech. Model. Mechanobiol.* **7,** 477–486 (2008).
13. Lu, J. & Zhao, X. Pointwise Identification of Elastic Properties in Nonlinear Hyperelastic Membranes—Part I: Theoretical and Computational Developments. *J. Appl. Mech.* **76** (2009).
14. Goldberg, M. A. A Linearized Large Deformation Analysis for Rotationally Symmetric Membranes. *J. Appl. Mech.* **32,** 444–445 (1965).
15. Lu, J. & Luo, Y. Solving membrane stress on deformed configuration using inverse elastostatic and forward penalty methods. *Comput. Methods Appl. Mech. Eng.* **308,** 134–150 (2016).
16. Romo, A., Badel, P., Duprey, A., Favre, J.-P. & Avril, S. In vitro analysis of localized aneurysm rupture. en. *J. Biomech.* **47,** 607–616 (2014).
17. Choi, P. T. & Lui, L. M. Fast Disk Conformal Parameterization of Simply-Connected Open Surfaces. *J. Sci. Comput.* **65,** 1065–1090 (2015).

18. Torres-Sánchez, A, Santos-Oliván, D & others. Approximation of tensor fields on surfaces of arbitrary topology based on local Monge parametrizations. *Journal of Computational* (2020).

19. Bonet, J. & Wood, R. D. *Nonlinear Continuum Mechanics for Finite Element Analysis* en (Cambridge University Press, 1997).

20. Schwarz, U. S. & Soiné, J. R. D. Traction force microscopy on soft elastic substrates: A guide to recent computational advances. en. *Biochim. Biophys. Acta* **1853,** 3095–3104 (2015).

21. Huang, Y. *et al.* Traction force microscopy with optimized regularization and automated Bayesian parameter selection for comparing cells. en. *Sci. Rep.* **9,** 539 (2019).

22. Tambe, D. T. *et al.* Collective cell guidance by cooperative intercellular forces. en. *Nat. Mater.* **10,** 469–475 (2011).

23. Tambe, D. T. *et al.* Monolayer stress microscopy: limitations, artifacts, and accuracy of recovered intercellular stresses. en. *PLoS One* **8,** e55172 (2013).

24. Stein, O., Jacobson, A., Wardetzky, M. & Grinspun, E. A Smoothness Energy without Boundary Distortion for Curved Surfaces. *ACM Trans. Graph.* **39,** 1–17 (2020).
